# *MembraneCurvature*: A Python tool to calculate membrane curvature from molecular dynamics simulations

**DOI:** 10.64898/2026.11.25.769279

**Authors:** Estefania Barreto-Ojeda, Valentina Corradi, D. Peter. Tieleman

## Abstract

Biological membranes are complex structural and functional assemblies that play central roles in many cellular processes. They are primarily composed of lipids and proteins, whose dynamics and interactions are commonly investigated at the molecular level using Molecular Dynamics (MD) simulations. Local membrane curvature, arising from changes in lipid composition and lipid–protein interactions, is an important property linked to lipid and protein sorting as well as protein mechanisms of action. Here, we present *MembraneCurvature*, a fast, user-friendly, and fully documented Python package designed to compute membrane curvature from MD simulations. Using selected lipid atoms as a reference, *MembraneCurvature* derives membrane surfaces and calculates molecular descriptors such as mean and Gaussian curvature. The package is built on top of the MDAnalysis library, a versatile framework compatible with simulations obtained from different software, force fields and membranes of simple or complex compositions. To demonstrate its application, we analyze MD simulations of a model plasma membrane generated using the Martini force field, both in the presence and absence of the phospholipid efflux ABC transporter ABCA1. In this example, *MembraneCurvature* identifies persistent, protein-induced membrane curvature that depends on the protein’s conformational state and that is leaflet-specific.

## 1. INTRODUCTION

Cell membranes are functional and structural assemblies involved in several biological processes, including cell trafficking, signaling, regulation, and cell architecture, and are considered main building blocks in living organisms.^1, 2^ They are composed primarily of a lipid matrix and associated proteins, whose structure and function are closely linked. The lipid matrix is arranged in a double layer of lipids, with their hydrophobic tails pointing towards the centre of the bilayer, and their hydrophilic headgroups pointing outwards. This arrangement confers fluidity and elasticity to the membrane and forms a barrier that many molecules cannot cross. The presence of membrane proteins provides additional properties such as selective permeability, allowing certain molecules or ions to cross it, depending on their size, shape, and charge; and curvature, where membrane bending is the result of lipids and protein distribution and protein shape.^3–8^ Lipid contribution to membrane curvature arises from an asymmetric lipid composition between leaflets, combined with the different shape and size of the lipid headgroups compared to the acyl tails.^4, 5^ Peripheral or transmembrane proteins can sense, induce or stabilize membrane curvature through various mechanisms, by inserting specific motifs or domains in the membrane, and/or by adopting a particular shape, as shown for example by the superfamily of the BAR proteins^9^ or by transmembrane proteins like the ATP-binding cassette transporter BmrA^10^ and the voltage-gated potassium channel KvAP.^11^ Thus, in biological membranes, membrane deformation and local membrane curvature are the results of the complex interplay between lipids and proteins, their different structure and shape, and preferable interactions, governing sorting mechanisms.^4, 5^ However, there are limited molecular details on how membrane curvature is affected by changes in lipids composition, protein adsorption, insertion or other events like protein-protein interactions.

Molecular dynamics (MD) simulations are a powerful tool to explore and analyze complex systems and have been widely used to characterize structure, dynamics and properties of biological membranes systems.^12–15^ Membrane curvature profiles can be derived from MD simulations using tools of differential geometry, combined with elastic theories developed since the 1970s.^16–18^ Each of the two monolayers of a membrane is described as a differentiable surface in the 3-dimensional Euclidean space, on which the principal curvatures, *c*1 and *c*2, are defined at every point as the maximum and minimum values of the normal curvature. To obtain curvature information, the mean (H = ½(*c*_1_ + *c*_2_), in Å^-1^) and Gaussian (K = (*c*_1_ *c*_2_), in Å^-2^) curvature metrics are calculated.^19^ The mean curvature provides information about the direction and magnitude of bending of the membrane patch taken into account: H > 0 values indicate convex (valleys) regions in the membrane, in which the surface curves upwards; H < 0 values are associated with concave (peaks) regions, in which the surface curves downwards; while H = 0 values represent either flat regions or symmetric saddles of the membrane. The Gaussian curvature describes the local shape of the membrane patch, independently of its orientation, where the surface corresponds to a saddle-like surface if K < 0; to a surface curved in at most one direction, such as a cylinder or a plane, if K = 0; and to a sphere-like surface if K > 0. A membrane patch is therefore locally flat only where H and K are both zero. Among the available tools that extract different features from simulation systems related to membrane shape, for example thickness, area per lipid, and lipid order parameter (GridMAT-MD,^20^ Membrainy,^21^ FATSlim,^22^ APL@Voro,^23^ LiPyphilic,^24^ MOSAICS^25^), only a few allow for calculation of membrane curvature: `g_lomepro`^26^ was developed to work specifically as a GROMACS analysis tool, CATpie^27^ can derive curvature, area, and thickness from MD simulations using a Fourier-based approach, while MemCurv^28^ was designed with particular applications in mind to study membrane remodeling by the FAM134B receptor. MemSurfer^29^ is a more general tool that leverages Delaunay triangulations to derive membrane surfaces, which can in turn be used to obtain thickness, area per lipid, lipid distribution and membrane curvature information from MD simulations. MembraneAnalysis.jl^30^ is a recent analysis tool developed in Julia that calculates bending modulus, spontaneous curvature and other properties like thickness of the membrane and correlations between them, and other tools are dedicated to highly curved and asymmetric membranes.^31, 32^ In all of these tools, however, the representation of the membrane surface and the treatment of temporal averaging in curvature calculations are coupled in their implementation. As a result, users cannot separate the contribution of the time-averaged membrane shape from that of the fluctuations around it, even though both carry different physical information, and in the case of Gaussian curvature, do not commute. In addition, Python is one of the most popular programming languages in scientific computing, and despite available tools, there is a need for an easy-to-use, well-documented, and regularly maintained Python-based tool to calculate membrane curvature from MD simulations.

Here, we introduce *MembraneCurvature*, a Python-based open-source tool dedicated to the calculation of mean and Gaussian curvature from MD simulations of membranes and membrane protein systems. The tool has the advantage of being built on top of the popular MDAnalysis library for analysis of MD simulations^33, 34^ and it is registered as an MDAKit in the MDAnalysis Toolkits registry,^35^ which supports FAIR principles (Findable, Accessible, Interoperable, Reusable).^36^ It works with both atomistic and coarse-grained (CG) simulation systems and can be installed on different operating systems. We present a detailed explanation of its algorithm, a description of its computational features, and illustrate its application using CG MD simulations of the human ABC transporter ABCA1 embedded in a plasma membrane model, and of the membrane alone. Finally, we discuss challenges and limitations of the simulation setup and the tool when analyzing curvature. *MembraneCurvature*is available at https://github.com/MDAnalysis/membrane-curvature.

## 2. METHODS

### 2.1 The MembraneCurvature tool

#### Overview

*MembraneCurvature* is an object-oriented, force-field agnostic tool to calculate membrane curvature from MD simulations. The tool is written in Python 3 and requires only two input files: A coordinate file and a trajectory file. Overall, the workflow used by *MembraneCurvature* includes five main steps (Figure 1). First, the user identifies the lipid atoms of reference, which define the monolayers of interest to derive the curvature profiles from (Figure 1, Step 1). Then, using the dimension of the simulation box, *MembraneCurvature* derives a surface from the selected reference group using one of three methods: The Fourier method (default) or one of the two binning methods, namely binning and binning nearest (Figure 1, Step 2 and 3). In the Fourier method, a truncated two-dimensional Fourier series is fitted to the atomic heights at each frame using linear least squares. In the binning methods, the surface is instead constructed on a grid at each frame: the height at each bin is defined as the mean z-coordinate of the reference atoms assigned to it (binning) or, alternatively, as the z coordinate of the reference atom nearest to it (binning nearest) in the *x*, *y* plane. The partial derivatives required to calculate the mean and Gaussian curvature for each frame of the trajectory are then obtained according to the chosen method (Figure 1, Step 4): For the binning methods, the derivatives are estimated numerically from the discrete height field and grid spacing, while for the Fourier method, they are evaluated analytically from the fitted Fourier series. Curvature can then be calculated either from the surfaces derived at every frame (Figure 1, Step 4A) or once, from the average surface obtained after the per-frame surface calculation (Figure 1, Step 4B). *MembraneCurvature* outputs the results in the form of surface, mean and Gaussian curvature arrays (Figure 1, Step 5).

**Figure 1.**
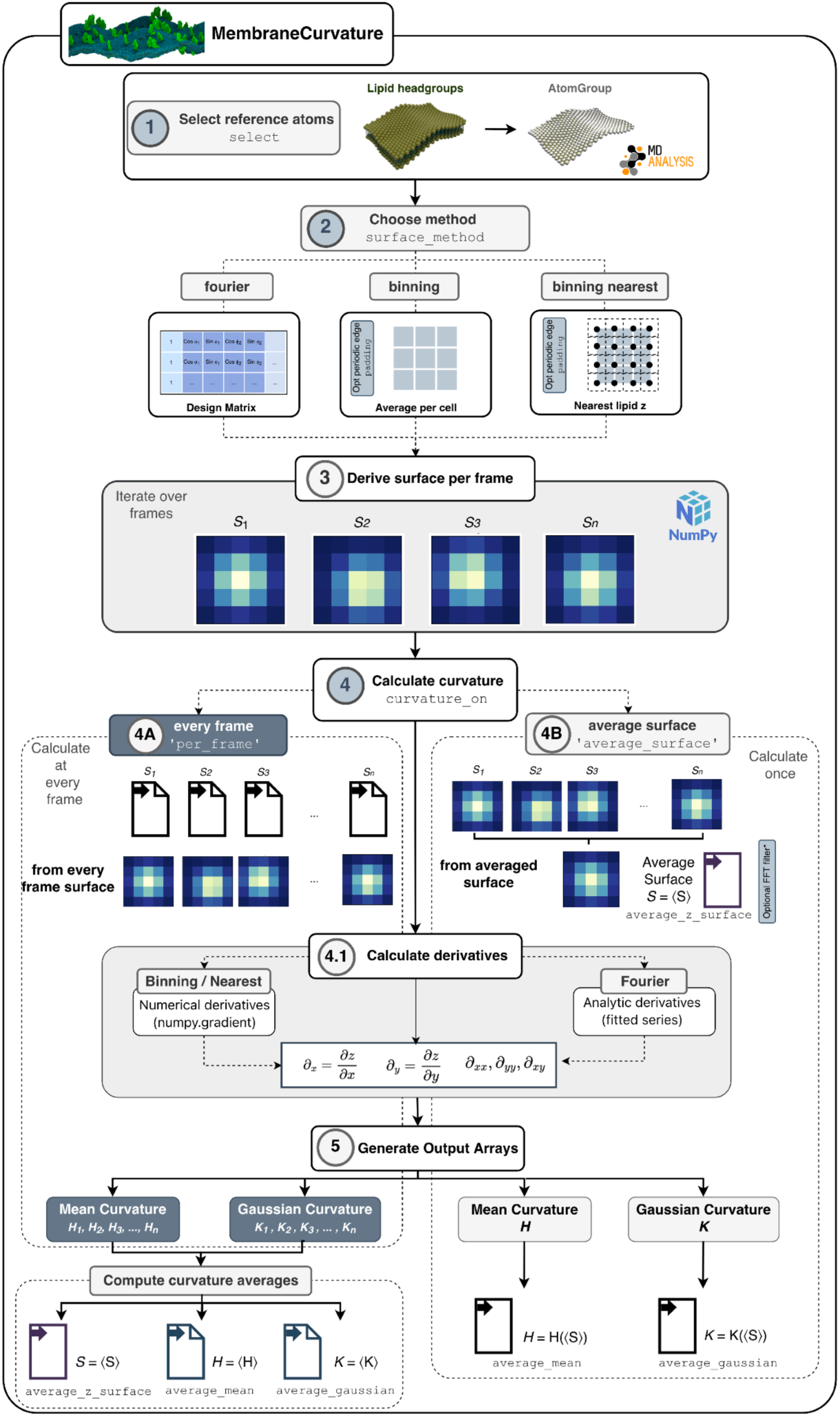
Overview of *MembraneCurvature* workflow. *MembraneCurvature* calculates mean and Gaussian curvature of surfaces derived from selected atom groups of reference using grid-based binning methods and a Fourier method.

#### Software engineering

*MembraneCurvature* has been developed and maintained under the MDAnalysis GitHub organization, where the full development history, issue tracker, and releases are available. Hosting within the organization ensures that the repository, its history, and maintainer access persist independently of any individual contributor account. *MembraneCurvature* is a free, open-source software package under the GNU General Public License, version 3 (GPLv3).

The design principles of *MembraneCurvature* can be classified into three main categories: testability, modularity, and versatility.

#### Testability

*MembraneCurvature* was developed using the test-driven and mutation testing practices in software development. Unit test coverage was increased progressively during development, reaching 100% code coverage. This ensures that every line of code is exercised by the test suite. *MembraneCurvature* is tested in three operating systems: MacOS, Linux and Windows.

#### Modularity

The functionality of *MembraneCurvature* is separated into independent and interchangeable modules corresponding to different stages of the analysis pipeline: grid construction and surface derivation for the binning methods, least-squares surface fitting for the Fourier method, periodic padding, FFT filtering, and curvature calculation. Input and parameter validation is handled by dedicated modules that check the reference selection, the box geometry, and the method-specific parameters before the analysis run and raise informative errors when a requested combination of settings is not supported. Building on these modules, *MembraneCurvature* uses the MDAnalysis AnalysisBase class as a building block to run multi-frame and average-over-frames analyses, deriving surfaces and their respective values of mean and Gaussian curvature. The modular code of *MembraneCurvature* is a key feature for code maintenance, improvement, and debugging.

#### Versatility

*MembraneCurvature* performs curvature calculations for both atomistic and coarse-grained representations of membrane systems. Because *MembraneCurvature* uses the MDAnalysis library as its main engine, it can be used with input structures and trajectory files from a wide variety of biomolecular simulation software supported by MDAnalysis, including for example, GROMACS,^37^ CHARMM,^38^ NAMD,^39^ Amber,^40^ and LAMMPS.^41^ In addition, *MembraneCurvature*’s Application Programming Interface (API) is simple and intuitive and allows the user to calculate and extract curvature results with a few lines of code.

#### Algorithm details

The algorithm implemented in *MembraneCurvature* includes three different methods (two binning methods and the Fourier method) to derive the surface from a selected group of atoms to calculate curvature. Both approaches generate a height field *Z_i_*_j_ ≈ *f*(*x_i_*, *y*_j_) on a regular *N_x_* × *N_y_* grid, together with the corresponding first and second partial derivatives of *f*. Given the partial derivatives of the height field *∂*_x_, *∂*_y_, *∂*_xx_, *∂*_yy_, *∂*_xy_ evaluated at each grid point, the mean curvature (*H*) and Gaussian curvature (*K*) in the Monge gauge framework are defined as:

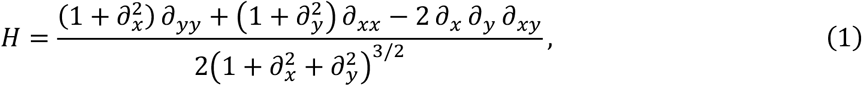

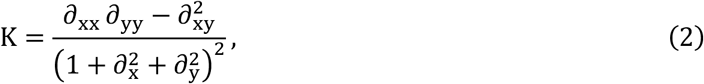

where 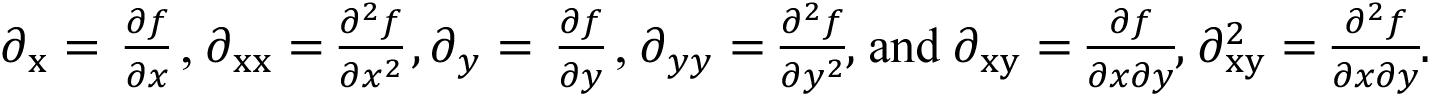 Below we describe in detail how each approach works according to the steps in Figure 1.

Step 1: Atoms of reference

*MembraneCurvature* derives one surface for a user-defined *AtomGroup*. In biological membranes, atoms (or beads in the case of Martini CG simulations) from lipid headgroups are typically selected as reference atoms for the upper and lower leaflet independently. The corresponding *AtomGroups* must be provided by the user as input using selection strings that follow standard MDAnalysis syntax.

Step 2 and 3: Choice of methods and surface calculation

*MembraneCurvature* can derive the surface corresponding to the reference atoms using grid-based binning methods (*i.e.*, binning and binning nearest) or the Fourier method (default).

#### The binning method (surface_method=’binning’)

After defining the reference atoms in an *AtomGroup*, a given simulation box of size *L_x_*, *L_y_* is partitioned into an *N_x_* × *N_y_* grid with bin widths *Δx* = *L_x_*/*N_x_*, *Δy* = *L_y_*/*N_y_*. *MembraneCurvature* automatically detects the size of the simulation box from the first frame of the trajectory, although these can be set explicitly by the user (parameters: `x_range`, `y_range`), as can the number of bins *N_x_* × *N_y_*. Each atom *α* of the reference *AtomGroup* with coordinates (*x_α_*, *y_α_*, *z_α_*) is mapped to a bin (*l*, *m*) given by:

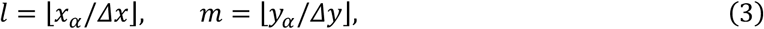

with *l* = 0,…,*N_x_* − 1 and *m* = 0,…,*N_y_* − 1.

For this method, coordinate wrapping is enabled by default (parameter: `wrap=True`), so that at each frame, atoms that fall outside the primary cell are wrapped back into it before populating the grid. Atoms that still map outside the valid index range are skipped, and a UserWarning reporting the number of skipped atoms is issued. The height assigned to each bin (*l*, *m*) is the arithmetic mean of the *z* coordinates of all the atoms assigned to the bin:

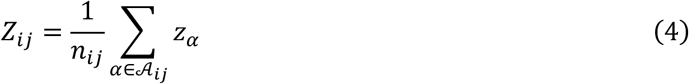

where A_ij_ is the set of atoms assigned to the bin (*l*, *m*) and *n_lm_*is the number of atoms in that bin. Unpopulated bins in the grid are left undefined and excluded from curvature averages. For every frame of the trajectory, the collection of all *Z* values across the grid defines the membrane surface *S*.

#### The binning nearest method (surface_method=’binning_nearest’)

The binning approach described above has the downside of finding potential empty bins for surface calculation, which propagate into the curvature calculation, resulting in regions of undefined curvature. To overcome this, we have implemented the binning nearest method, which differs in how the *z* coordinate of the chosen atoms of reference are assigned to the bins: The binning nearest method, similarly to the `g_lomepro` tool,^26^ builds the grid to derive a surface by assigning to each grid point the *z* coordinate of the nearest atom of reference in the *x*, *y* plane with periodic boundary conditions and with grid points sitting at bin corners. The resulting height field is then used to calculate curvature. This method differs from the `surface_method=’binning’` approach, as it evaluates the surface at bin corners and not at bin centers. Because the distance search already accounts for periodicity, coordinate wrapping does not apply and is disabled for this method.

For both binning methods, two additional options are available: Periodic padding and an FFT filter. Periodic padding: The grid-based binning methods do not explicitly account for the periodicity of the simulation box when calculating surface derivatives. To reduce artifacts at the box edges, particularly those arising in the calculation of second derivatives and Gaussian curvature, the option `padding=True` can be used. When enabled, *MembraneCurvature* expands the primary grid by a periodic buffer on each side. The resulting padded grid contains the original bins as well as additional bins populated using periodically wrapped height values. Mean and Gaussian curvature are then evaluated over the padded height field, after which the buffer is removed so that the returned curvature arrays correspond to the original primary grid. By default, `edge_pad_bins=2` when `padding=True`. This value is sufficient for the finite-difference calculation used to obtain the surface derivatives, which accesses the first neighboring bins on either side. Larger values of `edge_pad_bins` are accepted but provide no additional information and only increase computational cost.

A brick-wall FFT filter is also available for the binning methods through the `fft_filter` parameter. After the height fields have been computed for all the frames, *MembraneCurvature* calculates the time-averaged height field. If FFT filtering is enabled, this averaged surface is filtered once in reciprocal space by applying a brick-wall filter, in which Fourier modes with wavevector magnitudes (wavenumbers) outside the specified passband are set to zero before transforming the filtered surface back to real space. Individual per-frame surfaces are not FFT-filtered. These passband bounds (*q_lo_*_3_, *q*_ℎ*i*5ℎ_) determine the range of spatial scales retained in the membrane height field: Small values correspond to longer-wavelength, broader undulations, whereas large values correspond to shorter-wavelength, finer-scale undulations. The shortest wavelength that can be represented by the grid is limited by the Nyquist wavenumbers, which depend on the grid spacing in the *x*, *y* directions. The Nyquist wavenumbers along the *x*, *y* directions are *π*/Δ*x*, *π*/Δ*y*, respectively. For the isotropic radial filter used by *MembraneCurvature*, a single conservative Nyquist wavenumber is defined as the smaller of these two values: *q_Nyq_* = *min*(*π*/Δ*x*, *π*/Δ*y*). By default, `fft_filter=None`, and no FFT filtering is applied. With `fft_filter=’auto’`, a low-pass filter with a passband (*q_lo_*_3_, *q*_ℎ*i*5ℎ_) = (0, 0.5*q_Nyq_*) is applied. This provides a conservative cutoff, although the optimal upper cutoff may depend on the system and grid resolution. Alternatively, users can specify the bounds explicitly as `fft_filter={‘q’:(q_low, q_high)}`, in rad/Å. The FFT filter can also be combined with periodic padding.

#### The Fourier method

The Fourier method is the default method in *MembraneCurvature*. This method is related to Fourier surface modeling^42^ and molecular Fourier shape descriptors.^43^ However, as described below, it is not a direct implementation of either approach. Instead, the least-squares fit of the height field is specifically adapted to the *AtomGroup* of reference.

The membrane height field is represented as a real-valued, doubly periodic Fourier series truncated at mode indices *M* (in *x*) and *N* (in *y*). The basis functions are periodic over the simulation box, with periods *L_x_* and *L_y_*, to ensure that the fitted surface is consistent with periodic boundary conditions:

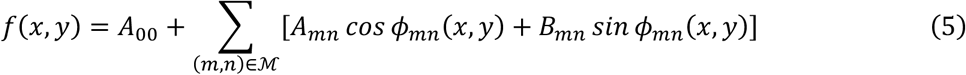

where *A*_00_ is the constant Fourier coefficient, corresponding to the spatially averaged membrane height (*i.e.*, mean height coefficient); *A_mn_* and *B_mn_* are the cosine and sine amplitudes of mode (*m*, *n*); and ℳ is the set of retained modes defined below. The phase associated with mode (*m*, *n*) is:

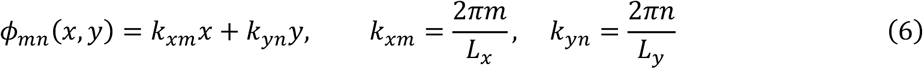

The unknown parameters are the mean term *A*_00_ and the cosine/sine amplitudes *A_mn_*, *B_mn_*. Because the height field is real-valued, the modes (*m*, *n*) and (−*m*, −*n*) are not independent so that *A*_−*m*,−*n*_ = *A_mn_* and *B*_−*m*,−*n*_ = *B_mn_*, and both modes contribute the same real basis functions. As retaining both would duplicate parameters, only a non-redundant subset of Fourier modes is kept, thereby reducing the number of fitted parameters and the computational cost of the least-squares solution. Specifically, the mode set ℳ is defined as:

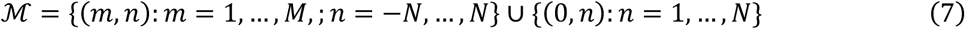

with the *A*_00_ coefficient stored separately. Since this set contains *M*(2*N* + 1) + *N* modes, then the total number of scalar fitting parameters is:

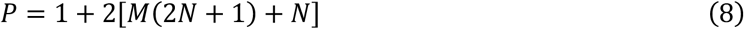

Consequently, the atom selection (*AtomGroup*) must contain at least *P* atoms. If it contains fewer than *P*, *MembraneCurvature* raises a ValueError and reports the largest mode index the selection can support. Before evaluating the Fourier basis functions, the *x* and *y* coordinates of each atom are shifted and wrapped into the fundamental periodic domain, [0, *L_x_*), ‘0, *L_y_*), according to:

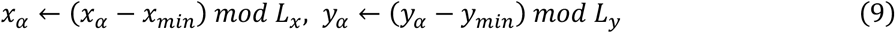

where *x_min_* and *L_x_* are the lower bound and the width of the domain along *x*, and similarly for *y*. By default, *MembraneCurvature* sets *x_min_* = *y_min_* = 0, and *L_x_*, *L_y_* from the simulation box dimensions. This transformation maps all atoms to equivalent positions within a single periodic unit cell while preserving the original heights *z_α_*. To assemble the design matrix **Φ**, the wrapped coordinates (*x*’*_a_*,*y’_a_*) are used to evaluate the Fourier basis functions at the position of each atom, building **Φ** to contain one row per atom. The first column of the design matrix **Φ** consists entirely of ones and corresponds to the mean height coefficient *A*_00_:

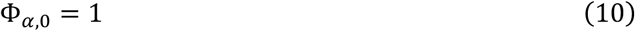

while each retained mode contributes one cosine and one sine basis function in adjacent columns:

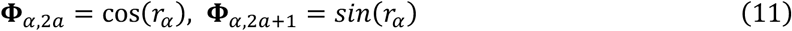

where (*r_α_*) = *k_xm_x_α_* + *k_yn_y_α_* and wavevectors are defined as in equation (6), and *a* is the index of the column where *a* ∈ [0, *K*) and *K* = *M*(2*N* + 1) + *N*, the number of non-redundant modes.

Once the design matrix is constructed, together with the fitting parameter **Θ**, a linear least squares system is formed and given by:

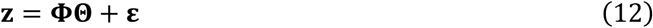

where ***z*** is the vector of heights of the *N_atoms_* in the reference AtomGroup, **Θ** is the parameter vector that contains the *P* unknown Fourier coefficients, **Φ** is the design matrix of dimensions N_atoms_ × *P*, and *ε* is the residual vector of length *N_atoms_*. As the model is linear in **Θ**, estimation of the coefficients reduces to an ordinary least squares problem:

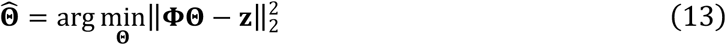

*MembraneCurvature* solves this system by Singular Value Decomposition (SVD), implemented via the `numpy.linalg.svd` routine in the NumPy library.^44^ After decomposing **Φ**, singular values below a relative threshold that satisfies *σ* ≤ *r_cond_* · *σ_max_* are treated as numerically zero, where *σ_max_* is the largest singular value of **Φ** and *r_cond_* is a relative cutoff. The effective rank of **Φ** is the number of singular values that exceed this threshold, and it indicates how many independent directions in the coefficient space are constrained by the heights of the reference atoms. When *r*_cond_ is not specified, *MembraneCurvature* sets the value as: *r_cond_* = *ε* · max(*N_atoms_*, *P*), where *ε* is float64 machine precision. This truncation improves numerical stability when the design matrix is rank-deficient. The solution of the linear system returns the fitted coefficient vector **Θ** = [*A*_00_, *A_m_*_1_*_n_*_1_, *B_m_*_1_*_n_*_1_,…,*A_m_*_K_*_n_*_K_, *B_m_*_K_*_n_*_K_]^T^, ordered consistently with the modes list ℳ, as in equation (7). Once the coefficient vector **Θ** is known, the fitted height field is evaluated analytically on a regular bin center mesh of *N_x_* × *N_y_* points:

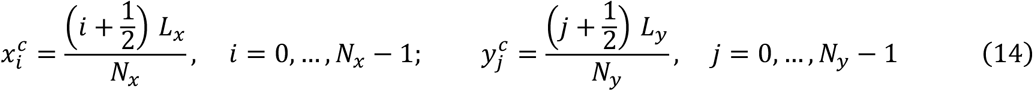

generating the array ***Z*** ∈ ℝ*^Nx^*^×*Ny*^ with entries:

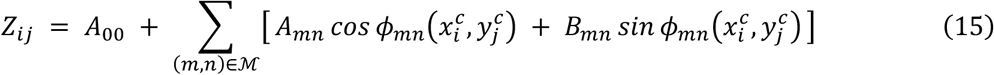

obtained by direct summation of the Fourier Series (equation 5) rather than by a fast Fourier transform. We note that the surface is a continuous function of (*x*, *y*) and the grid in this case is used solely for its (*N_x_* × *N_y_*) resolution, and it does not affect the fitted coefficients but only the sampling of the reconstructed surface. This grid resolution is the support on which the analytical partial derivatives are evaluated and mean and Gaussian curvature will be reported.

Independently from the method of choice to derive the surfaces from the atoms of reference, *MembraneCurvature* computes a surface for each frame of the input trajectory.

Step 4 and 5: Curvature calculation and output arrays

#### Binning methods

The partial derivatives required to obtain mean and Gaussian curvature (eq. 1 and 2) are estimated numerically from the discrete height field *Z* using finite differences:

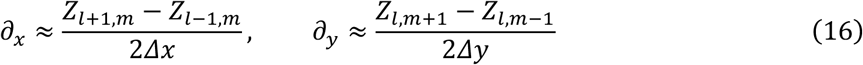

calculated using the NumPy package for scientific computing,^44^ with physical spacing Δ*x* and Δy. The second-order derivatives *∂*_x*x*_, *∂*_y*y*_, *∂*_xy_ are obtained by applying the same finite-difference operator a second time to the first derivatives array.

#### The Fourier method

Here, the partial derivatives are calculated analytically: For each mode (*m*, *n*) with phase *φ_mn_*(*x*, *y*) = *k_xm_ x* + *k_yn_y* and height contribution *z*(*φ*) = *A cos φ* + *B sin φ*, the chain rule gives:

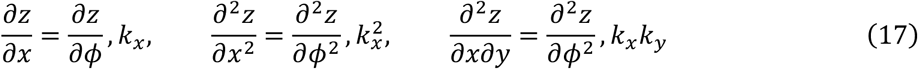

with *∂z*/*∂φ* = −*A sin φ* + *B cos φ* and *∂*^2^*z*/*∂φ*^2^ = −*A cos φ* − *B sin φ*. The five derivative arrays, *∂_x_*, *∂_y_*, *∂*_x*x*_, *∂*_y*y*_, *∂*_xy_, are accumulated by summing the contributions over all modes in ℳ and evaluating the result on the bin-centre mesh. Mean and Gaussian curvature are then computed from these arrays using the Monge-gauge formulas (equations 1 and 2).

#### Temporal averaging

For all methods, mean and Gaussian curvature can be calculated as average curvature maps evaluated from (i) the per-frame curvature maps (parameter: `curvature_on=’per_frame’`) or (ii) directly on the time-averaged surface height field (parameter: `curvature_on=’average_surface’`). By default, the mean and Gaussian curvature maps are averaged over all frames. For backward compatibility with older versions of *MembraneCurvature*, passing `curvature_on=None` (default) is equivalent to `curvature_on=’per_frame’`. For the binning methods, when using the FFT filter, if `curvature_on` is not provided, enabling `fft_filter` selects `curvature_on=’average_surface’`.

#### Outputs

For all methods, the results are stored as Numpy arrays. The per-frame surface and curvature maps are returned in the attributes `results.z_surface`, `results.mean` and `results.gaussian,` and their averages over the trajectory in `results.average_z_surface`, `results.average_mean` and `results.average_gaussian.` Mean and Gaussian curvature are reported in Å^-1^and Å^-2^, respectively.

#### Dependencies and installation

*MembraneCurvature* has only two main dependencies, MDAnalysis^33, 34^ (>=2.4.0) and NumPy,^44^ (>=1.23.0), and requires Python >=3.11. Keeping its own dependency declaration minimal simplifies long-term maintenance by delegating trajectory handling to MDAnalysis. The latest available release can be installed on the user’s operating system via the Python Package Index (PyPI) `pip`:^45^

`pip install membrane-curvature`

or via conda:^46^

conda install-c conda-forge membrane-curvature

or with uv:

uv pip install membrane-curvature

#### Documentation

Comprehensive documentation for *MembraneCurvature* is available at: https://mdanalysismembrane-curvature.readthedocs.io/en/latest/index.html, and the complete API reference at: https://mdanalysismembrane-curvature.readthedocs.io/en/latest/api.html.

The documentation includes:

- An explanation of the algorithm
- Usage examples
- Visualization options
- Tutorials for atomistic and coarse-grained simulation systems

### 2.2 Application of *MembraneCurvature*

#### Simulation systems

The simulation systems included in this work were generated using the CG Martini 3 force field.^47, 48^. All simulation steps, from system setup to production run, were performed with GROMACS^37^, v. 2024.4.

##### Systems Setup

###### Membrane only system

We used a plasma membrane model (namely, PM8) consisting of 8 lipid types asymmetrically distributed between upper and lower leaflet, as previously described^47^ (Table S1). The simulation system was built using INSANE,^49^ specifically its newest version available from the Martini 3 Lipidome^47^ set of tools (https://github.com/Martini-Force-Field-Initiative/M3-Lipid-Parameters/tree/main/tools/insane_M3_lipids_new.py), including water and a 0.15M NaCl. *ABCA1 systems.* We used PDB IDs 7TBY and 7BTW for the open and closed conformation, respectively, of ABCA1.^50^ The initial atomistic structures were processed through CHARMM-GUI^51^ to rebuild the residues 1788-1798 missing loop for the open conformation, and the residues 1467-1471 and 1835-1840 missing loops for the closed state. For the open state, we used the resulting structure as input to setup the corresponding Martini system. For the closed state, we performed additional atomistic steps, as initial tests revealed some instability in CG simulation tests. Specifically, the atomistic structure, after insertion in a POPC membrane via CHARMM-GUI,^52, 53^ was subjected to two rounds of energy minimization using CHARMM36m^54^ and CHARMM36^55^ forcefields for protein and lipids, respectively. The first energy minimization round initially included position restraints on the protein backbone and side chain atoms (force constant of 1000 and 20 700 kJ mol^-1^ nm^-2^, respectively), and on lipid atoms (force constant of 100 700 kJ mol^-1^ nm^-2^ for the phosphate atom and for the dihedral restraints). For the second round, no position restraints were included. For both rounds of energy minimization, the criteria for convergence were set at 50000 steps or a maximum force smaller than 10 kJ mol^-1^nm^-1^. The output of the second round of energy minimization was used as input for the Martini 3 setup of the closed state.

Each of the resulting ABCA1 structures were converted to their corresponding coarse-grained model for Martini 3 using `martinize2` (from vermouth version 0.15.0),^56^ choosing to generate one elastic network for the entire protein, using the default 700 kJ mol^-1^ nm^-2^ force constant and an upper cutoff limit of 0.9 nm for the elastic bonds, and selecting the default protein model available in Martini 3.^48^ Each protein conformation (*i. e.* ABCA1 open, ABCA1 closed) was then embedded in the PM8 membrane using INSANE as described for the membrane model system (Table S1).

##### Systems’ simulations

The three CG systems (*i. e.*, PM8, ABCA1 open, ABCA1 closed) were energy minimized by applying the steepest descent algorithm using, for convergence criteria, 5000 steps or a maximum force smaller than 10 kJ mol^-1^nm^-1^. This was followed by a 10 ns equilibration, which used the Berendsen thermostat and barostat,^57^ with a time constant of 1.0 and 5 ps for temperature and pressure coupling, respectively. Finally, each system was simulated for 30 μs, switching to the v-rescale thermostat^58^ and the c-rescale barostat,^59^ with time constant of 1.0 and 4 ps for temperature and pressure coupling, respectively, setting the temperature of the system at 310 K and the pressure at 1 bar. All the simulation parameters used for GROMACS correspond to those recommended for Martini 3 as provided by the latest Martini 3 Lipidome work.^47^

##### Simulation trajectories post-processing

The 30 μs-long trajectory of the PM8 system was post-processed with the GROMACS `gmx trjconv` tool using the following periodic boundary condition options: `-pbc nojump`, followed by `-pbc whole` and `-pbc res-ur compact` to make molecules whole and placed them back into the simulation box. For the ABCA1 open and closed trajectories, we additionally centered the protein in the simulation box `(-pbc res-ur compact-center`) and removed rotations and translations in the x and y directions `(-fit rotxy+transxy`). From these post-processed trajectories, we extracted 5 μs-long intervals, as well as 2 μs-long intervals for the PM8 system using `gmx trjconv`. Each interval was then further processed with MDAnalysis^33, 34^ (v. 2.9) so that the center of mass (COM) of the bilayer was centered at 0. For this, lipids that translocate between leaflets in the protein systems were excluded from the calculation of the COM.

##### Membrane curvature calculation

We used *MembraneCurvature* to calculate the mean and Gaussian curvature over 5 μs-long time windows of each simulation system. For all the calculations, the surface of each leaflet was derived using the PO4 beads of phospholipids as the *AtomGroup* of reference. Cholesterol was excluded due to its ability to flip flop. In addition, in the ABCA1 open and closed systems, we found 13 and 1, respectively, phospholipids that translocated between leaflets during the simulations. Such lipids were excluded from the surface calculation, while the PO4 beads of lipids entering the open cavity of ABCA1 without fully translocating were kept. No lipid translocations were detected for the membrane only systems. From these leaflet surfaces, maps of mean and Gaussian curvatures were then generated using binning methods and the Fourier method. As we processed the trajectory as described above with GROMACS tools, wrapping was disabled for these analyses (parameter: `wrap=None`). For the Fourier method, we used the default number of modes (*M* = 2, *N* = 2). For the binning methods, we tested different settings and bin sizes, and for the final figures we set the number of bins in the *x* and *y* dimensions to 10, resulting in unit cells of approximately 24-25 Å.

Standard deviation maps for surface, mean and Gaussian curvature were obtained for the 5 μs intervals of each simulation system, when the Fourier and the binning nearest method with periodic padding were applied. The surface, mean and Gaussian curvature maps from all intervals were stacked, and the standard deviation was calculated at each grid point across the time intervals. Standard deviations were calculated independently for the upper and lower leaflet.

All analyses were performed with *MembraneCurvature* 2.0.0, MDAnalysis 2.10 and python 3.14.5.

#### Figures

Figures for surfaces, mean and Gaussian were generated with the `pyplot` module of the Matplotlib library,^60^ using the `imshow` function and a bilinear interpolation method. Molecular representations of the ABCA1 system were generated with VMD.^61^

#### Computational performance

To determine the computational cost of *MembraneCurvature* calculations, we performed per-frame measurements on an Apple M4 Max (arm64) running macOS, with Python 3.14.5, NumPy 2.2.6 built against the Accelerate BLAS and LAPACK backend, MDAnalysis 2.10.0 and *MembraneCurvature* 2.0.0. All calculations used a 10 × 10 grid for the binning methods, and the Fourier method used the default number of modes (*M* = 2, *N* = 2). The reference *AtomGroup* is the PO4 beads of the lower leaflet of each system. Each reported value is the median of five repeats after one warm-up run and includes reading of the trajectory.

## 3. RESULTS

To illustrate the use of *MembraneCurvature*, we performed CG MD simulations of a model plasma membrane (namely, PM8)^47^ in the presence and absence of the human ATP-binding cassette transporter A1, in both the open, nucleotide free and the closed, nucleotide-bound state (here referred to as ABCA1 open and ABCA1 closed systems).^50^ ABCA1 plays a key role in the generation of high-density lipoprotein (HDL), by loading cholesterol and lipids such as phosphatidylcholine and phosphatidylserine into a lipid acceptor apolipoprotein A-I (apoA-I).^62, 63^ The structure of ABCA1 consists of two Nucleotide Binding Domains (NBDs), two transmembrane domains (TMD), each of six transmembrane helices (TMH), and two large extracellular domains (ECDs), required for substrate and apoA-I binding (Figure 2). In the nucleotide-free open state, the two TMDs form a cavity that is open towards the extracellular side, with a smaller opening on the intracellular side. This asymmetry shapes the cavity differently on each side of the membrane and allows lipids to bridge across the two TMDs (Figure 2, upper panels). In the closed state (Figure 2, lower panels), the TMDs are tightly bound, following the NBDs dimerization in the intracellular side, with no accessible central cavity.

**Figure 2.**
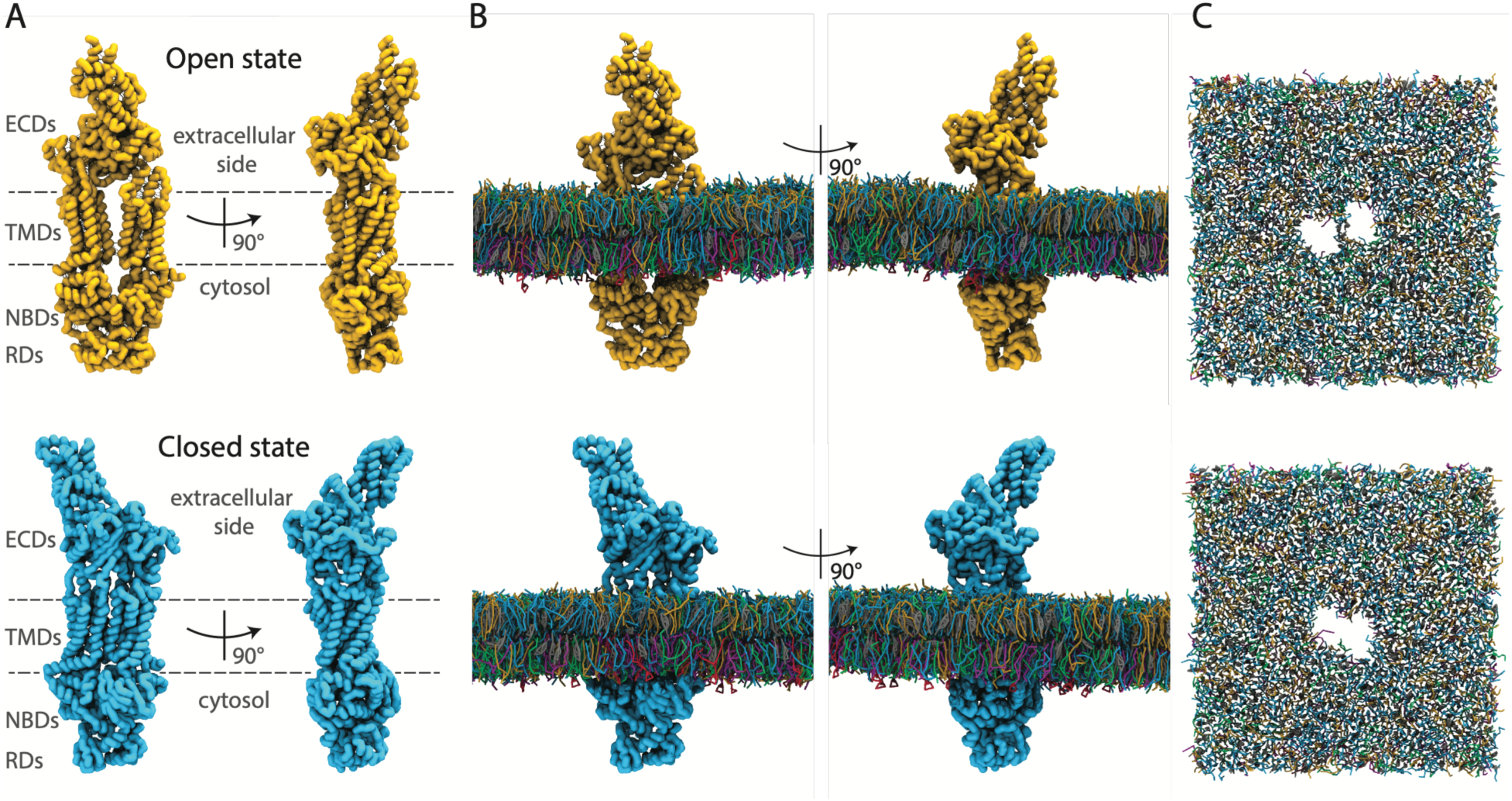
The ABCA1 transporter simulation systems. (A) CG structure of ABCA1 in the open (upper panel, PDB ID: 7TBY)^50^ and closed (lower panel, PDB ID 7TBW)^50^ state, shown as yellow and cyan surface, respectively. (B) The open (upper panel) and closed (lower panel) states of ABCA1 are shown embedded in the PM8 plasma membrane model at time 0 of the corresponding production runs. Water and ions are not shown for clarity. (C) View from the extracellular side of the two systems. The protein is not shown to highlight the different footprint of the two conformational states on the membrane. Lipids are shown as sticks and colored according to their headgroup type: PE, green; PC, cyan; Cholesterol: gray; SSM, dark yellow; SAP6, red; PS, magenta.

Each system (PM8, ABCA1 open and closed) was simulated for 30 μs, and curvature was calculated over 5 μs intervals, using the PO4 beads of the phospholipids as the *AtomGroup* of reference for each leaflet. First, we show how the surface methods and the optional settings implemented in *MembraneCurvature* affect the corresponding curvature maps of the upper leaflet of the PM8 system and the two conformational states of ABCA1, using the last 5 μs of each simulation. Then, curvature calculations are shown as a function of time.

### Surface method comparisons

The results of mean and Gaussian curvature for PM8 are shown in Figure 3 below. While the profile for the surface and mean curvature is largely consistent across the different surface methods and settings, Gaussian curvature shows a strong dependence on the method used, particularly near the edges of the box. This is because while mean curvature is linear in the second derivatives of the surface, Gaussian curvature, on the other hand, depends on products of second derivatives and on the square of mixed derivatives. For the binning methods, at the grid boundary, the finite differences become one-sided, and this amplifies the mixed derivative term and biases Gaussian curvature towards negative values. Periodic padding supplies the missing neighbors, removing the edge artifacts. In this regard, the Fourier method is unaffected, since its basis is periodic and its derivatives are analytic.

**Figure 3.**
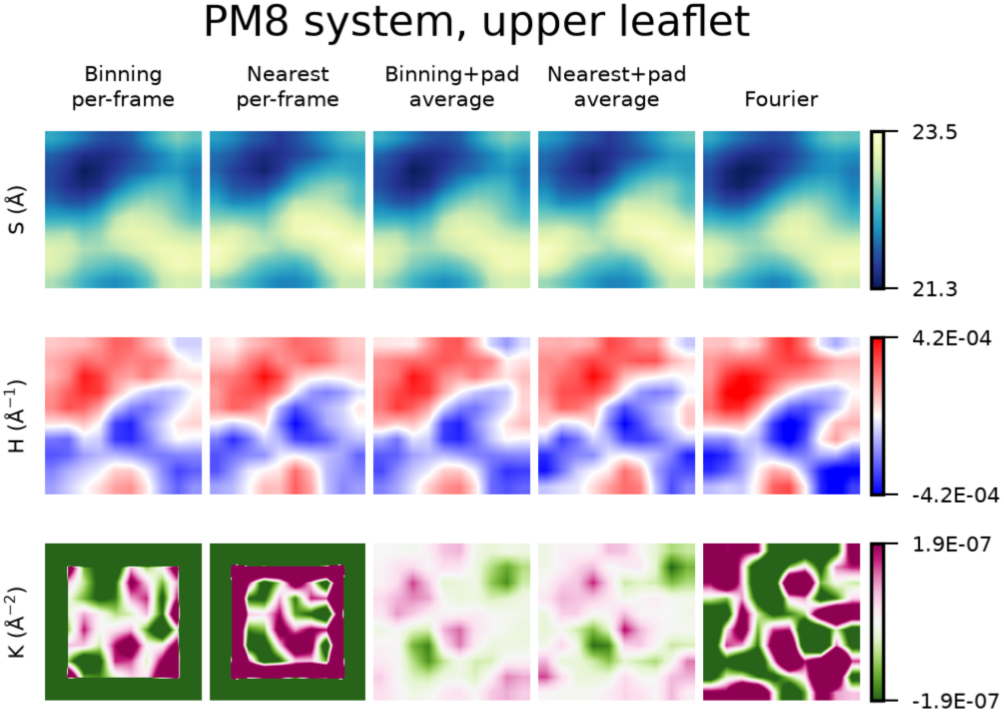
Effect of the surface methods, periodic padding and temporal averaging options on the curvature maps of the PM8 system. The last 5 μs of the upper leaflet of the PM8 simulation system were used to showcase the application of different settings in *MembraneCurvature*. From left to right: Binned and nearest-neighbour surfaces with curvature computed per frame; the same two methods with periodic padding and curvature computed on the time-averaged surface; and a Fourier-series fit (m = n = 2). Exact parameters are given in Table S2.

Figure 3 also highlights how Gaussian curvature depends which surface curvature is calculated from, set by the `curvature_on` parameter: The Fourier method and the binning method with curvature derived from per-frame maps (`curvature_on=’per_frame’`) produce consistently higher-magnitude Gaussian-curvature features than the binning nearest method when curvature is calculated on the averaged surface (see for example: ‘Nearest+pad average’ in Figure 3). This difference reflects the order in which averaging and differentiation are performed: With the `curvature_on=’average_surface’` parameter, mean and Gaussian curvature are calculated from the average surface array, not from the per-frame arrays, thus per-frame fluctuations are averaged before differentiation. This difference is expected to be more pronounced for Gaussian than for mean curvature: In the small-gradient regime relevant to these membranes, mean curvature is essentially a linear functional of the height field, (*H* ≈ ½(*∂_xx_* + *∂_yy_*))B. Then averaging the per-frame curvature maps and calculating curvature on the time-averaged surface give nearly the same result. Gaussian curvature, in contrast, is quadratic, (*K* ≈ (*∂_xx_∂_yy_* − *∂*^2^))B. Hence, calculating curvature on the averaged surface removes these fluctuation terms, whereas averaging per-frame maps retains them. Overall, for PM8 the results are consistent with those obtained from the `g_lomepro` tool^26^ (Figure S1).

The same algorithms and settings comparison were performed for the upper leaflet of the ABCA1 systems (Figure 4). One difference relative to `g_lomepro` results is detected for the ABCA1 open state, where the upper leaflet from *MembraneCurvature* shows a larger separation between the region of positive curvature near the protein (Figure 4, Figure S1).

**Figure 4.**
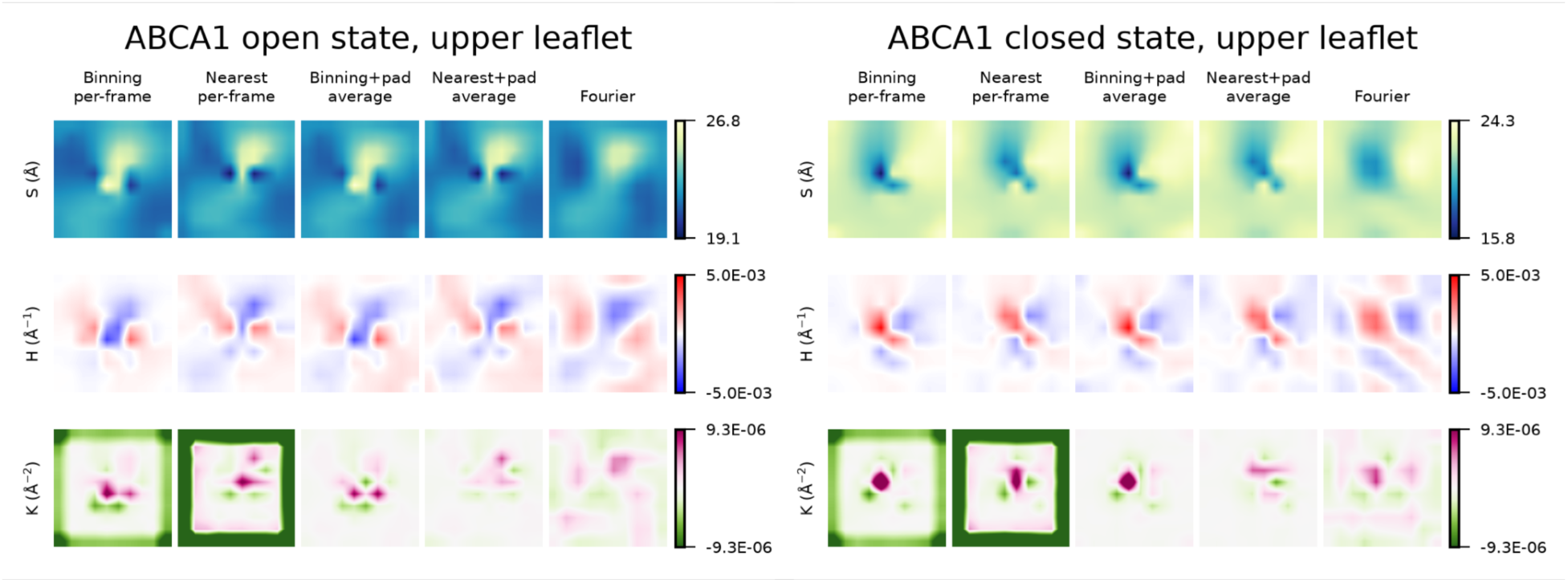
Effect of the surface methods, periodic padding and temporal averaging options on the curvature maps of the ABCA1 systems. The comparison is shown only for the upper leaflet over the last 5 μs of each simulation system. The algorithms and settings used corresponds to those described in Figure 3 for the PM8 system.

As mentioned above, the binning methods depend on the number of bins selected to construct the grid. As shown for the last 5 μs of the upper leaflet of the ABCA1 closed state simulation, a lower number of bins (*e.g.*, 6) results in coarser curvature profiles, and loss of spatial details (Figure S2). Increasing the number of bins decreases the size of individual grid cells and therefore increases the spatial resolution; however, this can also lead to insufficient sampling within individual cells (Figure S2). This is particularly relevant in the presence of membrane proteins, which occupy regions of the membrane and can reduce or exclude lipid sampling in the corresponding grid cells. As a result, some grid cells may contain no sampled height values, resulting in undefined surface values. As curvature is calculated from spatial derivatives of the surface, these gaps can also affect curvature estimates in neighboring cells. It is worth nothing, though, that the occurrence of empty grid cells also depends on the length of the trajectory analyzed, as longer trajectories provide more opportunities for lipids to sample individual cells: Here, we show an application of *MembraneCurvature* using a 5 μs window, but grid cells that remain unsampled at a given number of bins over this 5 μs window may contain sampled values when a longer time window is considered. In contrast, the Fourier method represents the surface as a continuous Fourier expansion and calculates its derivatives analytically, so localized regions with reduced or absent lipid sampling, such as those occupied by membrane proteins, do not directly result in missing curvature values. However, as described below in the “curvature as a function of time interval” section, the choice of the time window used to construct the average surface remains important for both binning and Fourier methods.

For the binning methods, *MembraneCurvature* also implements a FFT filter that can be applied in the reciprocal to retain only large-scale undulations of the membrane, when mean and Gaussian curvature is too noisy. This is achieved by using the following parameters: `fft_filter=’auto’, curvature_on=’average_surface’`, and by including periodic padding to remove the artifacts due to the intrinsic lack of periodicity with the binning methods (`padding=True`). The results are comparable with those obtained from `g_lomepro`, which also has the option of applying a FFT filter in both the real and reciprocal space (Figure S3).

### Curvature maps as a function of the simulation time

We calculated mean and Gaussian curvature using 5 μs-long time windows spanning the 30 μs of each simulation system and compared the results with the membrane only system. The results were obtained using the Fourier method, with the default number of modes and with the binning nearest method, using periodic padding and by calculating curvature using the `curvature_on=’average_surface’` parameter.

#### Membrane surface

For the simulation systems with ABCA1, membrane shape remains relatively stable over time, using 5 μs-long intervals (Figure S4). In both conformational states, the lower leaflet surface near the TMDs shows an upward, hill-like deformation. This feature, however, is more prominent for the ABCA1 open state, where it is also coupled with a similar deformation profile in the upper leaflet (Figure S4). In contrast, over the same time scale, the PM8 system shows greater variability, with surface profiles derived from 5 μs-long time windows that differ from one another (Figure S5).

Combined, these results indicate that the presence of ABCA1 induces and stabilizes membrane deformations, leading to more persistent structural features that correlate with the protein’s conformational state. In comparison, the membrane alone (PM8) produces transient, localized surface features that dissipate rapidly, relative to the timescale of the deformations of the ABCA1 systems.

#### Mean and Gaussian Curvature

For the ABCA1 systems, the 2D maps for mean curvature reveal persistent structural features, retrieved using both the Fourier method and the binning nearest method with periodic padding (Figure 5A-B and S6A-B). In the open state, the region immediately surrounding the cavity, which corresponds to higher z coordinate values of the PO4 beads in both upper and lower leaflet, is characterized by negative mean curvature that propagates diagonally, in the lower leaflet, flanked by regions of positive mean curvature (Figure 5A, S6A). For the ABCA1 closed state, the lower leaflet exhibits a similar profile to that of the open state (Figure 5B, S6B). In contrast, the upper leaflet showed a strong region of positive curvature, disrupting the negative curvature pattern detected in the ABCA1 open state (Figure 5B, S6B). While the results described here for the Fourier method are reproduced by the binning nearest method, when using periodic padding, we note that the difference in intensity in mean and Gaussian curvature between upper and lower leaflet is less strong with the binning method. Overall, the curvature patterns of each conformational state persist over all the 5 μs-long intervals spanning the 30 μs simulations. Compared to the PM8 simulation (Figure S7), the curvature profile induced by the protein shows a greater magnitude. However, mean curvature values remain on the order of 10^-3^ _Å_-1.

**Figure 5.**
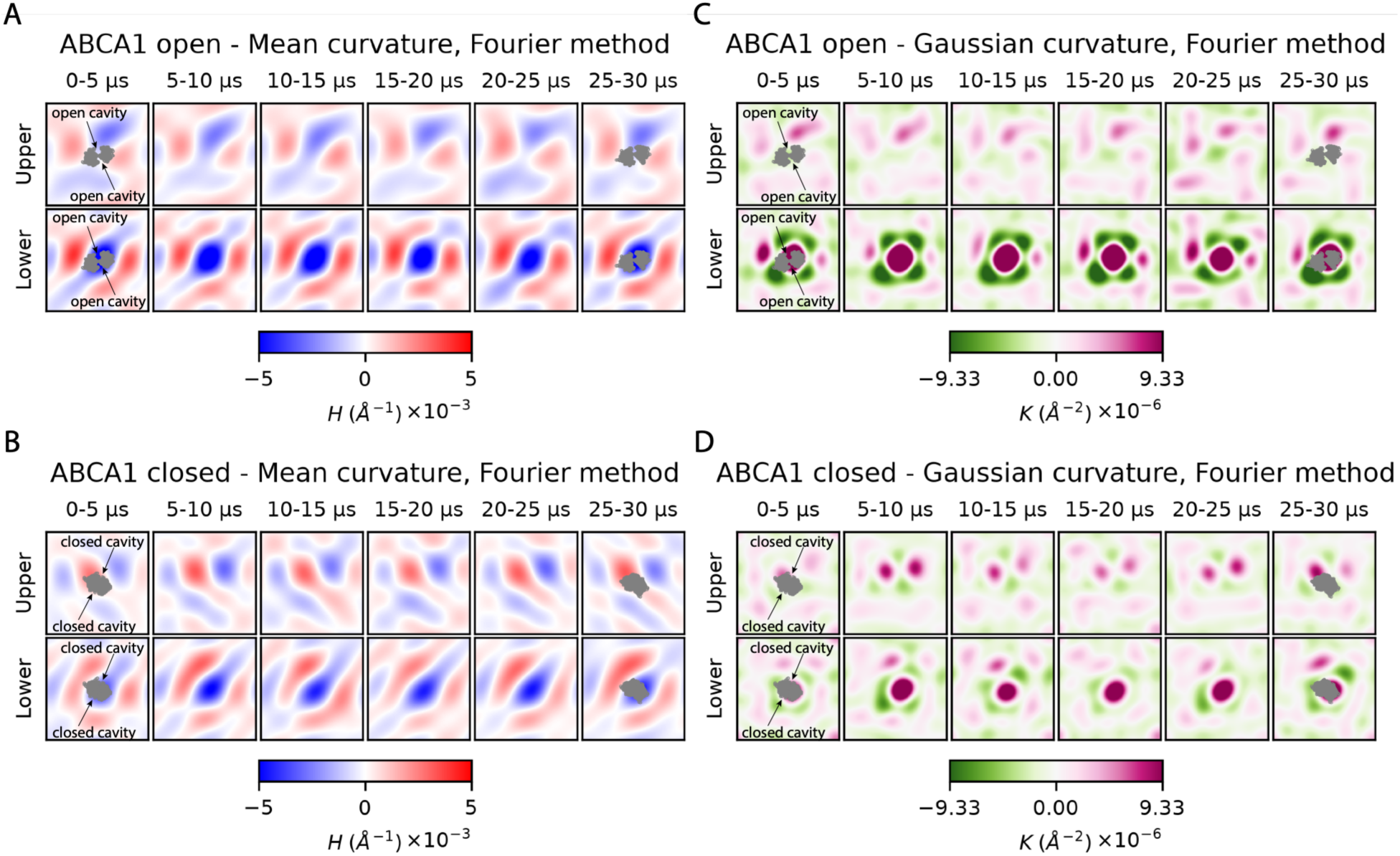
2D maps of mean and Gaussian curvature from the ABCA1 open and closed state systems obtained with the Fourier method. For the 0-5 and 25-30 μs intervals, the backbone of the protein TMDs at time 0 and 30 μs, respectively, was overlapped to the 2D maps.

For Gaussian curvature, both the open and closed state of ABCA1 (Figure 5C-D, S6C-D) produced distinct patterns of alternating positive, zero and negative values. In the lower leaflet, these patterns consist of a localized region of positive curvature (magenta) at the position of the TMDs, surrounded by a saddle-shaped region of negative curvature (green), which is more continuous and pronounced in the open state. In the upper leaflet, the profiles are different between open and closed state. As observed for mean curvature, the Gaussian curvature of the protein systems revealed more persistent features over the timescale and intervals considered for this analysis, compared to the PM8 simulation (Figure S7) but the scale of these perturbations remains small (on the order of 10^-6^ Å^-2^).

#### Standard deviation of surface, mean and Gaussian curvature

To assess the temporal variability at each spatial location of the surface, mean and Gaussian curvature derived from the 5 μs intervals above, we calculated the corresponding standard deviation maps for each leaflet for the Fourier method and the binning nearest method with periodic padding (Figure S8). Both methods show similar patterns, with larger regions of higher deviations in the lower leaflet for surface and mean curvature. For the ABCA1 systems, regions of higher variability are often located around the protein. However, these deviations are smaller than 1 Å, and the surface map of the PM8 system is within the same range. The range of deviations for mean curvature is within approximately 0-6 x10^-4^ Å^-1^ for the three systems. Gaussian curvature deviations are smaller for the PM8 system (ranging from approximately 0 to 7 x10^-7^ Å^-2^) compared with a 0 to 7 x10^-6^ Å^-2^ range for the ABCA1 systems.

### Surface and curvature maps as a function of the time interval

Surface and curvature profiles depend on the length and position of the time window used for the analysis, reflecting the range of timescales over which the membrane fluctuates: In protein-containing systems, as shown by the ABCA1 simulations above, protein-membrane interactions may stabilize local deformations, making some curvature features more persistent than those observed in protein-free membranes. To show the effect of the window length on curvature results, we also calculated the surface and curvature maps for the PM8 system using the Fourier method over 2 μs intervals (Figure S9-S10). Surface and curvature profiles differ not only from those obtained from 5 μs windows (Figure S5 and S7), but also between consecutive intervals. The standard deviation maps calculated from the 2 μs intervals show slightly higher values, reflecting the reduced temporal averaging in shorter windows and the larger contribution of short time-scale fluctuations (Figure S11). Therefore, the appropriate time window should be selected based on the system and the temporal or conformational fluctuations of interest.

### Computational performance

We measured the cost per frame of each surface method implemented in *MembraneCurvature*, for the simulation systems used in this work (Table 1). Binning is the least expensive method, at 0.60 to 0.63 ms per frame across the systems tested. Binning nearest requires between 1.1 and 1.2 times longer, and the Fourier method, using the default number of modes, requires approximately 1.7 times the cost of binning. The addition of the periodic padding changes the cost of both binning methods by less than 5%, and in some cases reduces it. The values reported in Table 1 include the reading of the trajectory, which accounts for a large fraction of the total in every case: For the PM8 system, the difference between the least and the most expensive method is 0.45 ms per frame, against a total of 1.08 ms per frame for the Fourier method. The cost in the Fourier method depends on the number of modes through the size of the design matrix, of dimensions *N_atoms_* × *P*. The analysis of 5 μs trajectory of the PM8 system, corresponding to 2501 frames, required between 1.6 and 2.9 s depending on the method.

**Table 1.** Cost per frame, in ms, of each surface method implemented in *MembraneCurvature*, using the curvature_on=’per frame’ parameter.

| System | Number of reference atoms | Number of frames | Binning | Nearest | Binning+pad | Nearest+pad | Fourier |
|---|---|---|---|---|---|---|---|
| PM8 | 912 | 2501 | 0.63 | 0.71 | 0.63 | 0.70 | 1.08 |
| ABCA1 open | 877 | 2501 | 0.60 | 0.74 | 0.62 | 0.74 | 1.09 |
| ABCA1 closed | 881 | 2501 | 0.63 | 0.78 | 0.64 | 0.76 | 1.11 |

## 4. DISCUSSION and CONCLUSIONS

We introduced *MembraneCurvature*, a MDAnalysis-based Python tool to calculate membrane curvature from atomistic and CG MD simulations of membrane and membrane-protein systems. *MembraneCurvature* is a MDAKit,^35^ which ensures that the tool is continuously tested against the latest development and stable versions of MDAnalysis, thereby maintaining compatibility with the evolving MDAnalysis ecosystem. This framework also enforces standardized software practices, including automated testing and validation, to ensure robustness, reproducibility, and long-term maintainability of the codebase.^35, 36^

In *MembraneCurvature* the derivation of the surface and temporal averaging of the curvature are controlled by independent and documented parameters, allowing users to run each option independently to evaluate their effect on the resulting curvature maps. The membrane surface can be derived using binning-based approaches and a Fourier-based approach (Table 2), and curvature can be evaluated either on each individual frame, and then averaged, or on a time-averaged surface (Table 3). A summary with limitations and advantages of each approach is presented in Tables 2 and 3.

**Table 2.** Surface derivation methods and boundary treatments implemented in *MembraneCurvature*.

| Surface derivation approach | Parameter | Limitations | Advantages |
|---|---|---|---|
| Binning | surface_method=’binning’ | Empty bins may occur; edge artifacts in curvature calculations | Fast |
| Binning nearest | surface_method=’binning_nearest’ | Surface maps are offset by half a bin relative to raw binning and Fourier methods; edge artifacts in curvature calculations | No empty bins |
| Periodic padded binning | surface_method=’binning’<br>OR<br>surface_method=’binning_nearest’<br>AND<br>padding=True | Higher computational cost; currently limited to orthorhombic simulation boxes | No edge artifacts by extending the surface periodically |
| Fourier | surface_method=’Fourier’ | Higher computational cost | Intrinsically periodic surface; analytic derivatives; no empty bins; no edge artifacts |

**Table 3.** Curvature calculation and temporal averaging implemented in *MembraneCurvature*.

| Curvature calculation approach | Parameter | Procedure | Limitations | Advantages |
|---|---|---|---|---|
| Average of per-frame curvature maps | curvature_on='per_frame' (or None) | A curvature map is calculated independently for each frame; the final curvature map is obtained by averaging the per-frame curvature maps over time | Noisier | Provides frame-resolved curvature information before averaging |
| Curvature of the time-averaged surface | curvature_on='average_surface' | The membrane surface is first averaged over time; a single curvature map is then calculated from the resulting time-averaged surface | Per-frame fluctuations are lost | Produces smoother maps and highlights the average membrane shape |
| FFT filter on the average surface for binning methods | surface_method=’binning’<br>OR<br>surface_method=’binning_nearest’<br>AND<br>padding=True<br>curvature_on='average_surface'<br>AND<br>fft_filter=’auto’ | A FFT filter is applied to the time-averaged surface before curvature is calculated | Might result in loss of information | Reduced high-frequency noise in the averaged surface |

We showed the application of *MembraneCurvatur*e to CG simulations of the open and closed state of the ABCA1 phospholipid efflux protein, embedded in a model plasma membrane, and of the model plasma membrane alone. Membrane bending induced by ABCA1 has been identified as a key feature in the lipid translocase mechanism as it provides anchoring for apoA-I,^62^ and atomistic simulations of ABCA1 have shown changes in membrane thickness and bending due to the presence of the protein.^64^ More recently, cryoEM data of ABCA1 in nanodiscs further confirmed the protein-induced curvature of the membrane.^65^ Here, with *MembraneCurvature*, we found that ABCA1 actively reshapes the membrane in its vicinity: In the upper leaflet, the membrane curvature near the protein is state-dependent, as reflected by the different mean curvature profiles of the two ABCA1 simulations (Figure 5A-B). In the lower leaflet, on the other hand, the membrane is perturbed in a similar way in both ABCA1 states as shown by the negative mean curvature region (Figure 5A-B), but can undergo more pronounced localized deformations in the open state, as indicated by the Gaussian curvature profiles (Figure 5C-D). While the Gaussian curvature remains on the same order of magnitude (10^-6^ Å^-2^) as in the lipids only system, the ABCA1 systems exhibit larger local magnitudes, within the timescale of the analysis. Thus, for the lipid composition and timescale of the simulations used here, there is a leaflet asymmetry in how ABCA1 states induce stress in the membrane, although it should be noted that the magnitude of Gaussian curvature remains smaller than that estimated in phenomena like virus-induced budding.^66^ *MembraneCurvature* results are consistent with the work of Sunidhi et al., where mean curvature in the region surrounding ABCA1, was derived from atomistic simulations using `g_lomepro`.^64^ While values of mean curvature are not reported in this work, membrane deformation is described surrounding the same region of the protein and with the same sign as the one detected by *MembraneCurvature*.

### Practical considerations when using *MembraneCurvature*

When calculating membrane curvature from MD simulations, different aspects of the simulation setup are important. Membrane undulations depend on the timescale of the simulations, in turn linked to the complexity of the system: Lipid composition, number of proteins, and system size as well as the level of approximation (atomistic or CG representation) can affect sampling of changes in membrane shape and corresponding curvature profiles. *MembraneCurvature* users can assess convergence of their simulations by plotting surfaces and curvature results derived from time intervals of different length and as a function of time. Of note, it has recently been reported that in GROMACS,^37^ one of the most widely used simulation software, the barostat, the frequency of updating the neighbour list and the choice of appropriate cutoffs can introduce unrealistic changes in membranes shape, especially for larger simulation systems,^67^ and should be carefully reviewed by users at the time of system setup.

The default Fourier method implemented in *MembraneCurvature* overcomes the limitations of grid-based binning methods, including defining the number of bins and adding periodic padding. The selection of the size of the unit cell is important, as the surfaces are derived from lipid head groups assigned to cells of a grid. We have shown here how the choice of number of bins can result in undefined values of mean and Gaussian curvature, in regions occupied by the protein, and in its immediate proximity. As described above, this is due to the definition of mean and Gaussian curvature as gradients calculated based on adjacent values. Therefore, the existence of a single, undefined result in the surface spreads to the corresponding values of mean and Gaussian curvature, translating in undefined cells in the curvature grids. This behavior is system-dependent and can be affected by the time scale of the simulation. *MembraneCurvature* users, if using binning methods, should test a range of number of bins and choose according to the size and complexity of their simulation systems. And while adding a periodic padding is optional, users are encouraged to enable it when using binning methods, to avoid artifacts near the edge of the box.

Finally, one advantage of using *MembraneCurvature* to analyze MD simulations is that the tool stores surface information in NumPy arrays. This enables users to utilize the arrays to calculate additional membrane descriptors, such as thickness, to derive correlations with mean and Gaussian curvature data, and to obtain standard deviations maps like those presented in this paper (Figure S8, S11) or averages derived from multiple replicas of a given system, which can lead to a better understanding of a protein mechanism of action.^68^

### Current limitations

The current implementation of *MembraneCurvature* supports planar bilayer patches, and curvature cannot be calculated for systems like vesicles, capsids or micelles. Future development could account for non-planar bilayers as well as for lipids that flip-flop between leaflets. In addition, the periodic padding for the binning methods is supported only for orthorhombic simulation boxes.

In conclusion, *MembraneCurvature* is a fully documented Python tool that adds to the current landscape of automated analysis tools for membrane simulation systems and easily integrates the calculation of membrane curvature with the widely used MDAnalysis library. Through curvature analyses, the tool will help users characterize the molecular details of how membrane bending is affected by proteins, lipid distribution, protein-protein, and lipid-protein interactions.

## ASSOCIATED CONTENT

### Supporting Information

The Supporting Information file includes figures S1-S11 and Tables S1-S2. The trajectory files will be made available in Zenodo upon acceptance of the manuscript.

### Author Contributions

EB-O and DPT: Conceptualization of the project; EB-O: *MembraneCurvature* Python tool development and implementation. EB-O and VC: Simulations and analyses. All authors contributed equally to the writing of the manuscript and have given approval to its final version.

### Funding Sources

EB-O was supported by Google Summer of Code.

This work was also supported by the Natural Sciences and Engineering Research Council (Canada) and the Canadian Institutes of Health Research. Further support came from the Canada Research Chairs program. Calculations were carried out on Digital Research Alliance of Canada infrastructures, funded by the Canada Foundation for Innovation and partners.

### Notes

The authors declare no competing financial interest.

## ACKNOWLEDGMENT

We would like to thank Dr. Oliver Beckstein and Dr. Tyler Reddy from the MDAnalysis community for their valuable contributions and expertise during the development of the *MembraneCurvature* repository.

## ABBREVIATIONS

MD: molecular dynamics;
CG: coarse-grained;
ABCA1: ATP-Binding Cassette transporter A1;
FAIR: findable, accessible, interoperable and reusable;
FFT: fast Fourier Transform.

## REFERENCES

(1) van Meer, G.; Voelker, D. R.; Feigenson, G. W. Membrane lipids: where they are and how they behave. Nat. Rev. Mol. Cell Biol. 2008, 9 (2), 112–124. DOI: 10.1038/nrm2330

(2) Cheng, X.; Smith, J. C. Biological Membrane Organization and Cellular Signaling. Chem. Rev. 2019, 119 (9), 5849–5880. DOI: 10.1021/acs.chemrev.8b00439

(3) Cooke, I. R.; Deserno, M. Coupling between lipid shape and membrane curvature. Biophys. J. 2006, 91 (2), 487–495. DOI: 10.1529/biophysj.105.078683

(4) Callan-Jones, A.; Sorre, B.; Bassereau, P. Curvature-driven lipid sorting in biomembranes. Cold Spring Harb. Perspect. Biol. 2011, 3 (2). DOI: 10.1101/cshperspect.a004648

(5) McMahon, H. T.; Boucrot, E. Membrane curvature at a glance. J. Cell Sci. 2015, 128 (6), 1065–1070. DOI: 10.1242/jcs.114454

(6) Zimmerberg, J.; Kozlov, M. M. How proteins produce cellular membrane curvature. Nat. Rev. Mol. Cell Bio. 2006, 7 (1), 9–19. DOI: 10.1038/nrm1784

(7) González, D. V.; Makowski, M.; Lillo, M. P.; Cao-Garcia, F. J.; Melo, M. N.; Almendro-Vedia, V. G.; López-Montero, I. Rotation of the c-Ring Promotes the Curvature Sorting of Monomeric ATP Synthases. Adv. Sci. 2023. DOI: 10.1002/advs.202301606

(8) Lipowsky, R. Remodeling of Membrane Shape and Topology by Curvature Elasticity and Membrane Tension. *Adv*. Biol. 2022, 6 (1). DOI: 10.1002/adbi.202101020

(9) Simunovic, M.; Voth, G. A.; Callan-Jones, A.; Bassereau, P. When Physics Takes Over: BAR Proteins and Membrane Curvature. Trends Cell Biol. 2015, 25 (12), 780–792. DOI: 10.1016/j.tcb.2015.09.005

(10) Fribourg, P. F.; Chami, M.; Sorzano, C. O. S.; Gubellini, F.; Marabini, R.; Marco, S.; Jault, J. M.; Levy, D. 3D Cryo-Electron Reconstruction of BmrA, a Bacterial Multidrug ABC Transporter in an Inward-Facing Conformation and in a Lipidic Environment. J. Mol. Biol. 2014, 426 (10), 2059–2069. DOI: 10.1016/j.jmb.2014.03.002

(11) Quemeneur, F.; Sigurdsson, J. K.; Renner, M.; Atzberger, P. J.; Bassereau, P.; Lacoste, D. Shape matters in protein mobility within membranes. Proc. Natl. Acad. Sci. U. S. A. 2014, 111 (14), 5083–5087. DOI: 10.1073/pnas.1321054111

(12) Corradi, V.; Sejdiu, B. I.; Mesa-Galloso, H.; Abdizadeh, H.; Noskov, S. Y.; Marrink, S. J.; Tieleman, D. P. Emerging Diversity in Lipid-Protein Interactions. Chem. Rev. 2019, 119 (9), 5775–5848. DOI: 10.1021/acs.chemrev.8b00451

(13) Enkavi, G.; Javanainen, M.; Kulig, W.; Rog, T.; Vattulainen, I. Multiscale Simulations of Biological Membranes: The Challenge To Understand Biological Phenomena in a Living Substance. Chem. Rev. 2019, 119 (9), 5607–5774. DOI: 10.1021/acs.chemrev.8b00538

(14) Marrink, S. J.; Corradi, V.; Souza, P. C. T.; Ingolfsson, H. I.; Tieleman, D. P.; Sansom, M. S. P. Computational Modeling of Realistic Cell Membranes. Chem. Rev. 2019, 119 (9), 6184–6226. DOI: 10.1021/acs.chemrev.8b00460

(15) Muller, M. P.; Jiang, T.; Sun, C.; Lihan, M. Y.; Pant, S.; Mahinthichaichan, P.; Trifan, A.; Tajkhorshid, E. Characterization of Lipid-Protein Interactions and Lipid-Mediated Modulation of Membrane Protein Function through Molecular Simulation. Chem. Rev. 2019, 119 (9), 6086–6161. DOI: 10.1021/acs.chemrev.8b00608

(16) Canham, P. B. The minimum energy of bending as a possible explanation of the biconcave shape of the human red blood cell. J. Theor. Biol. 1970, 26 (1), 61–81. DOI: 10.1016/S0022-5193(70)80032-7

(17) Helfrich, W. Elastic Properties of Lipid Bilayers - Theory and Possible Experiments. Z Naturforsch C 1973, *C 28* (11-1), 693-703. DOI: DOI 10.1515/znc-1973-11-1209

(18) Evans, E. A. Bending Resistance and Chemically-Induced Moments in Membrane Bilayers. Biophys. J. 1974, 14 (12), 923–931. DOI: Doi 10.1016/S0006-3495(74)85959-X

(19) Deserno, M. Fluid lipid membranes: from differential geometry to curvature stresses. Chem. Phys. Lipids 2015, 185, 11–45. DOI: 10.1016/j.chemphyslip.2014.05.001

(20) Allen, W. J.; Lemkul, J. A.; Bevan, D. R. GridMAT-MD: a grid-based membrane analysis tool for use with molecular dynamics. J. Comput. Chem. 2009, 30 (12), 1952–1958. DOI: 10.1002/jcc.21172

(21) Carr, M.; MacPhee, C. E. Membrainy: a ‘smart’, unified membrane analysis tool. Source Code Biol. Med. 2015, 10, 3. DOI: 10.1186/s13029-015-0033-7

(22) Buchoux, S. FATSLiM: a fast and robust software to analyze MD simulations of membranes. Bioinformatics 2017, 33 (1), 133–134. DOI: 10.1093/bioinformatics/btw563

(23) Lukat, G.; Kruger, J.; Sommer, B. APL@Voro: a Voronoi-based membrane analysis tool for GROMACS trajectories. J. Chem. Inf. Model. 2013, 53 (11), 2908–2925. DOI: 10.1021/ci400172g

(24) Smith, P.; Lorenz, C. D. LiPyphilic: A Python Toolkit for the Analysis of Lipid Membrane Simulations. J. Chem. Theory Comput. 2021, 17 (9), 5907–5919. DOI: 10.1021/acs.jctc.1c00447

(25) Bernhardt, N.; Faraldo-Gomez, J. D. MOSAICS: A software suite for analysis of membrane structure and dynamics in simulated trajectories. Biophys. J. 2023, 122 (11), 2023–2040. DOI: 10.1016/j.bpj.2022.11.005

(26) Gapsys, V.; de Groot, B. L.; Briones, R. Computational analysis of local membrane properties. J. Comput. Aided Mol. Des. 2013, 27 (10), 845–858. DOI: 10.1007/s10822-013-9684-0

(27) Hakami Zanjani, A. A. CATpie: A Python tool for quantifying membrane curvature, area, and geometrically faithful normal-line thickness from molecular dynamics simulations. BBA - Biomembranes 2026, 1868 (3), 184530. DOI: 10.1016/j.bbamem.2026.184530

(28) Bhaskara, R. M.; Grumati, P.; Garcia-Pardo, J.; Kalayil, S.; Covarrubias-Pinto, A.; Chen, W.; Kudryashev, M.; Dikic, I.; Hummer, G. Curvature induction and membrane remodeling by FAM134B reticulon homology domain assist selective ER-phagy. Nat. Commun. 2019, 10 (1), 2370. DOI: 10.1038/s41467-019-10345-3

(29) Bhatia, H.; Ingolfsson, H. I.; Carpenter, T. S.; Lightstone, F. C.; Bremer, P. T. MemSurfer: A Tool for Robust Computation and Characterization of Curved Membranes. J. Chem. Theory Comput. 2019, 15 (11), 6411–6421. DOI: 10.1021/acs.jctc.9b00453

(30) Hossein, A.; Sodt, A. j. MembraneAnalysis.jl: A Julia package for analyzing molecular dynamics simulations of lipid membranes. J. Open Source Softw. 2023, 8 (87), 5380. DOI: 10.21105/joss.05380

(31) Yesylevskyy, S. O.; Ramseyer, C. Determination of mean and Gaussian curvatures of highly curved asymmetric lipid bilayers: the case study of the influence of cholesterol on the membrane shape. Phys. Chem. Chem. Phys. 2014, 16 (32), 17052–17061. DOI: 10.1039/c4cp01544d

(32) Fabian, B.; Javanainen, M. Diffusion Analyses along Mean and Gaussian-Curved Membranes with CurD. J. Phys. Chem. Lett. 2024, 15 (11), 3214–3220. DOI: 10.1021/acs.jpclett.4c00338

(33) Michaud-Agrawal, N.; Denning, E. J.; Woolf, T. B.; Beckstein, O. MDAnalysis: a toolkit for the analysis of molecular dynamics simulations. J. Comput. Chem. 2011, 32 (10), 2319–2327. DOI: 10.1002/jcc.21787

(34) Gowers, R. J.; Linke, M.; Barnoud, J.; Reddy, T. J. E.; Melo, M. N.; Seyler, S. L.; Domanski, J.; Dotson, D. L.; Buchoux, S.; Kenney, I. M.;, et al. MDAnalysis: A Python package for the rapid analysis of molecular dynamics simulations. Proceedings of the 15th Python in Science Conference 2016, 98–105.

(35) Alibay, I.; Wang, L.; Naughton, F.; Kenney, I.; Barnoud, J.; Gowers, R. J.; Beckstein, O. MDAKits: A framework for FAIR-compliant molecular simulation analysis. In Proceedings of the 22nd Python in Science Conference (SCIPY 2023), Austin, TX, USA, 2023; Meghann Agarwal, Chris Calloway, Niederhut, D., Eds.; pp 76-84. DOI: 10.25080/gerudo-f2bc6f59-00a.

(36) Amaro, R. E.; Aqvist, J.; Bahar, I.; Battistini, F.; Bellaiche, A.; Beltran, D.; Biggin, P. C.; Bonomi, M.; Bowman, G. R.; Bryce, R. A.;, et al. The need to implement FAIR principles in biomolecular simulations. Nat. Methods 2025, 22 (4), 641–645. DOI: 10.1038/s41592-025-02635-0

(37) Abraham, M. J.; Murtola, T.; Schulz, R.; Páll, S.; Smith, J. C.; Hess, B.; Lindahl, E. GROMACS: High performance molecular simulations through multi-level parallelism from laptops to supercomputers. SoftwareX 2015, 1-2, 19-25. DOI: 10.1016/j.softx.2015.06.001

(38) Brooks, B. R.; Brooks, C. L., 3rd; Mackerell, A. D., Jr.; Nilsson, L.; Petrella, R. J.; Roux, B.; Won, Y.; Archontis, G.; Bartels, C.; Boresch, S.; et al. CHARMM: the biomolecular simulation program. J. Comput. Chem. 2009, 30 (10), 1545–1614. DOI: 10.1002/jcc.21287

(39) Phillips, J. C.; Hardy, D. J.; Maia, J. D. C.; Stone, J. E.; Ribeiro, J. V.; Bernardi, R. C.; Buch, R.; Fiorin, G.; Henin, J.; Jiang, W.;, et al. Scalable molecular dynamics on CPU and GPU architectures with NAMD. J. Chem. Phys. 2020, 153 (4), 044130. DOI: 10.1063/5.0014475

(40) Case, D. A.; Cheatham, T. E., 3rd; Darden, T.; Gohlke, H.; Luo, R.; Merz, K. M., Jr.; Onufriev, A.; Simmerling, C.; Wang, B.; Woods, R. J. The Amber biomolecular simulation programs. J. Comput. Chem. 2005, 26 (16), 1668–1688. DOI: 10.1002/jcc.20290

(41) Thompson, A. P.; Aktulga, H. M.; Berger, R.; Bolintineanu, D. S.; Brown, W. M.; Crozier, P. S.; in ‘t Veld, P. J.; Kohlmeyer, A.; Moore, S. G.; Nguyen, T. D.;, et al. LAMMPS - a flexible simulation tool for particle-based materials modeling at the atomic, meso, and continuum scales. Comp. Phys. Comm. 2022, 271, 108171. DOI: 10.1016/j.cpc.2021.108171

(42) Shen, L.; Kim, S.; Saykin, A. J. Fourier method for large scale surface modeling and registration. Comput. Graph. 2009, 33 (3), 299–311. DOI: 10.1016/j.cag.2009.03.002

(43) Leicester, S. E.; Finney, J. L.; Bywater, R. P. Description of molecular surface shape using Fourier descriptors. J. Mol. Graph. 1988, 6 (2), 104–108. DOI: 10.1016/0263-7855(88)85008-2

(44) Harris, C. R.; Millman, K. J.; van der Walt, S. J.; Gommers, R.; Virtanen, P.; Cournapeau, D.; Wieser, E.; Taylor, J.; Berg, S.; Smith, N. J.;, et al. Array programming with NumPy. Nature 2020, 585 (7825), 357–362. DOI: 10.1038/s41586-020-2649-2

(45) *Python Package Index - PyPI*; Python Software Foundation: https://pypi.org.

(46) Anaconda, Anaconda Software Distribution; https://anaconda.com.

(47) Pedersen, K. B.; Ingolfsson, H. I.; Ramirez-Echemendia, D. P.; Borges-Araujo, L.; Andreasen, M. D.; Empereur-Mot, C.; Melcr, J.; Ozturk, T. N.; Bennett, W. F. D.; Kjolbye, L. R.;, et al. The Martini 3 Lipidome: Expanded and Refined Parameters Improve Lipid Phase Behavior. ACS Cent. Sci. 2025, 11 (9), 1598–1610. DOI: 10.1021/acscentsci.5c00755

(48) Souza, P. C. T.; Alessandri, R.; Barnoud, J.; Thallmair, S.; Faustino, I.; Grunewald, F.; Patmanidis, I.; Abdizadeh, H.; Bruininks, B. M. H.; Wassenaar, T. A.;, et al. Martini 3: a general purpose force field for coarse-grained molecular dynamics. Nat. Methods 2021, 18 (4), 382–388. DOI: 10.1038/s41592-021-01098-3

(49) Wassenaar, T. A.; Ingolfsson, H. I.; Bockmann, R. A.; Tieleman, D. P.; Marrink, S. J. Computational Lipidomics with insane: A Versatile Tool for Generating Custom Membranes for Molecular Simulations. J. Chem. Theory Comput. 2015, 11 (5), 2144–2155. DOI: 10.1021/acs.jctc.5b00209

(50) Sun, Y.; Li, X. Cholesterol efflux mechanism revealed by structural analysis of human ABCA1 conformational states. *Nat*. Cardiovasc. Res. 2022, 1 (3), 238–245. DOI: 10.1038/s44161-022-00022-y

(51) Jo, S.; Kim, T.; Iyer, V. G.; Im, W. CHARMM-GUI: a web-based graphical user interface for CHARMM. J. Comput. Chem. 2008, 29 (11), 1859–1865. DOI: 10.1002/jcc.20945

(52) Wu, E. L.; Cheng, X.; Jo, S.; Rui, H.; Song, K. C.; Davila-Contreras, E. M.; Qi, Y.; Lee, J.; Monje-Galvan, V.; Venable, R. M.;, et al. CHARMM-GUI Membrane Builder toward realistic biological membrane simulations. J. Comput. Chem. 2014, 35 (27), 1997–2004. DOI: 10.1002/jcc.23702

(53) Feng, S.; Park, S.; Choi, Y. K.; Im, W. CHARMM-GUI Membrane Builder: Past, Current, and Future Developments and Applications. J. Chem. Theory Comput. 2023, 19 (8), 2161–2185. DOI: 10.1021/acs.jctc.2c01246

(54) Huang, J.; Rauscher, S.; Nawrocki, G.; Ran, T.; Feig, M.; de Groot, B. L.; Grubmuller, H.; MacKerell, A. D., Jr. CHARMM36m: an improved force field for folded and intrinsically disordered proteins. Nat. Methods 2017, 14 (1), 71–73. DOI: 10.1038/nmeth.4067

(55) Huang, J.; MacKerell, A. D., Jr. CHARMM36 all-atom additive protein force field: validation based on comparison to NMR data. J. Comput. Chem. 2013, 34 (25), 2135–2145. DOI: 10.1002/jcc.23354

(56) Kroon, P. C.; Grunewald, F.; Barnoud, J.; van Tilburg, M.; Brasnett, C.; Souza, P. C. T.; Wassenaar, T. A.; Marrink, S. J. Martinize2 and Vermouth provide a unified framework for molecular topology generation. eLife 2025, 12. DOI: 10.7554/eLife.90627

(57) Berendsen, H. J. C.; Postma, J. P. M.; Vangunsteren, W. F.; Dinola, A.; Haak, J. R. Molecular-Dynamics with Coupling to an External Bath. J. Chem. Phys. 1984, 81 (8), 3684–3690. DOI: 10.1063/1.448118

(58) Bussi, G.; Donadio, D.; Parrinello, M. Canonical sampling through velocity rescaling. J. Chem. Phys. 2007, 126 (1). DOI: 10.1063/1.2408420

(59) Bernetti, M.; Bussi, G. Pressure control using stochastic cell rescaling. J. Chem. Phys. 2020, 153 (11). DOI: 10.1063/5.0020514

(60) Hunter, J. D. Matplotlib: A 2D Graphics Environment. Comput. Sci. Eng. 2007, 9 (3), 90–95. DOI: 10.1109/MCSE.2007.55

(61) Humphrey, W.; Dalke, A.; Schulten, K. VMD: visual molecular dynamics. J. Mol. Graph. 1996, 14 (1), 33–38, 27-38. DOI: 10.1016/0263-7855(96)00018-5

(62) Phillips, M. C. Is ABCA1 a lipid transfer protein? J. Lipid Res. 2018, 59 (5), 749–763. DOI: 10.1194/jlr.R082313

(63) Quazi, F.; Molday, R. S. Differential phospholipid substrates and directional transport by ATP-binding cassette proteins ABCA1, ABCA7, and ABCA4 and disease-causing mutants. J. Biol. Chem. 2013, 288 (48), 34414–34426. DOI: 10.1074/jbc.M113.508812

(64) Sunidhi, S.; Sacher, S.; Atul; Garg, P.; Ray, A. Elucidating the Structural Features of ABCA1 in its Heterogeneous Membrane Environment. Front. Mol. Biosci. 2021, 8, 803078. DOI: 10.3389/fmolb.2021.803078

(65) Plummer-Medeiros, A. M.; Culbertson, A. T.; Morales-Perez, C. L.; Liao, M. Activity and Structural Dynamics of Human ABCA1 in a Lipid Membrane. J. Mol. Biol. 2023, 435 (8), 168038. DOI: 10.1016/j.jmb.2023.168038

(66) Schmidt, N. W.; Mishra, A.; Wang, J.; DeGrado, W. F.; Wong, G. C. Influenza virus A M2 protein generates negative Gaussian membrane curvature necessary for budding and scission. J. Am. Chem. Soc. 2013, 135 (37), 13710–13719. DOI: 10.1021/ja400146z

(67) Kim, H.; Fábián, B.; Hummer, G. Neighbor List Artifacts in Molecular Dynamics Simulations. J. Chem. Theory Comput. 2023, 19 (23), 8919–8929. DOI: 10.1021/acs.jctc.3c00777

(68) Barreto Ojeda, E. Interplay between membrane curvature and conformational states in ABC transporters. University of Calgary, Calgary, Canada, Retrieved from https://prism.ucalgary.ca., 2022.

