## Supporting Information for "*MembraneCurvature*: A Python tool to calculate membrane curvature from molecular dynamics simulations"

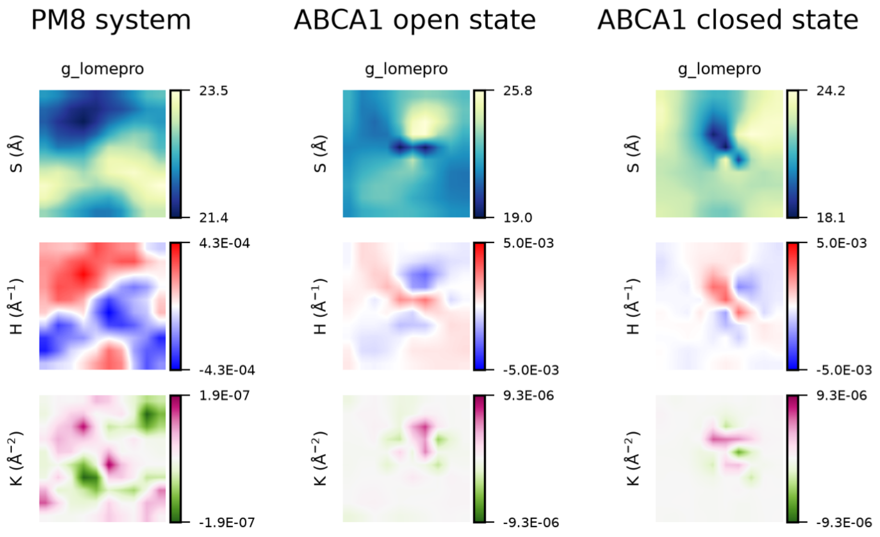


**Figure S1. Curvature profile of the PM8 and ABCA1 systems obtained with `g_lomepro`.**

The upper leaflet from the last 5 µs of the PM8 and ABCA1 simulations were also analyzed with the `g_lomepro` tool for comparison with *MembraneCurvature*. The `g_lomepro` default settings were used for the analysis.

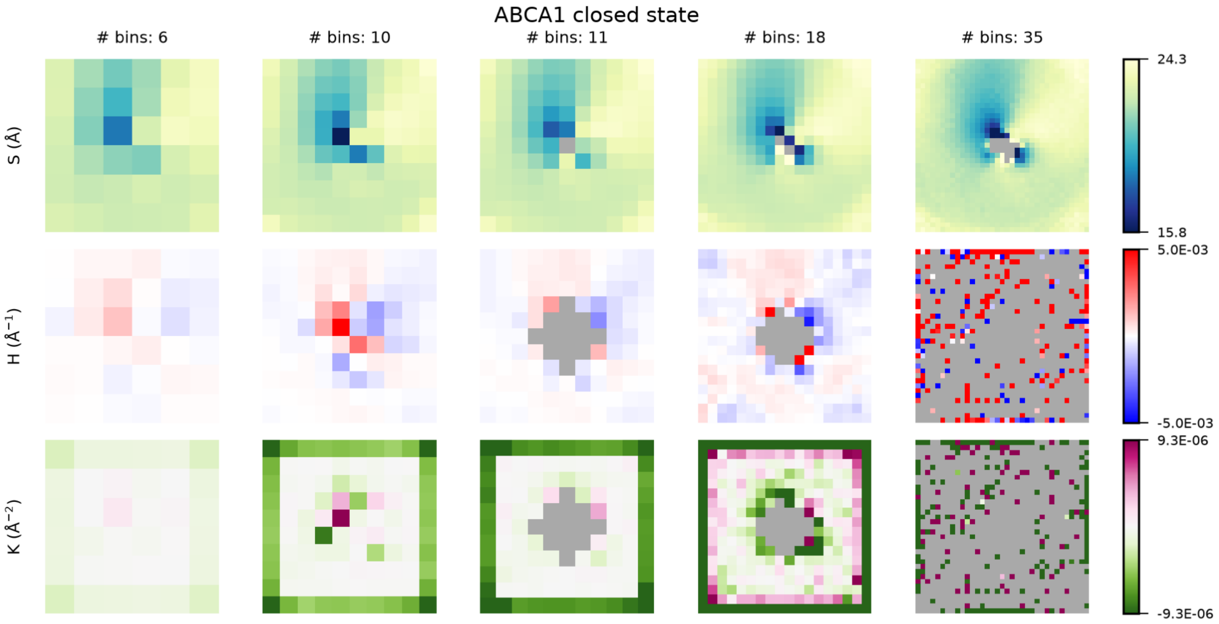


**Figure S2. ABCA1 closed state system: Impact of the number of bins in *MembraneCurvature* calculations.**

Undefined values in the surfaces used for curvature calculations result in undefined curvature regions. For the 25-20 µs time window, surface and curvatures maps were calculated for the upper leaflet of the ABCA1 closed state simulation using a different number of bins. The undefined values in maps are in dark gray. The 2D maps were obtained using `surface_method='binning'`, with no periodic padding, and were plotted without interpolation.

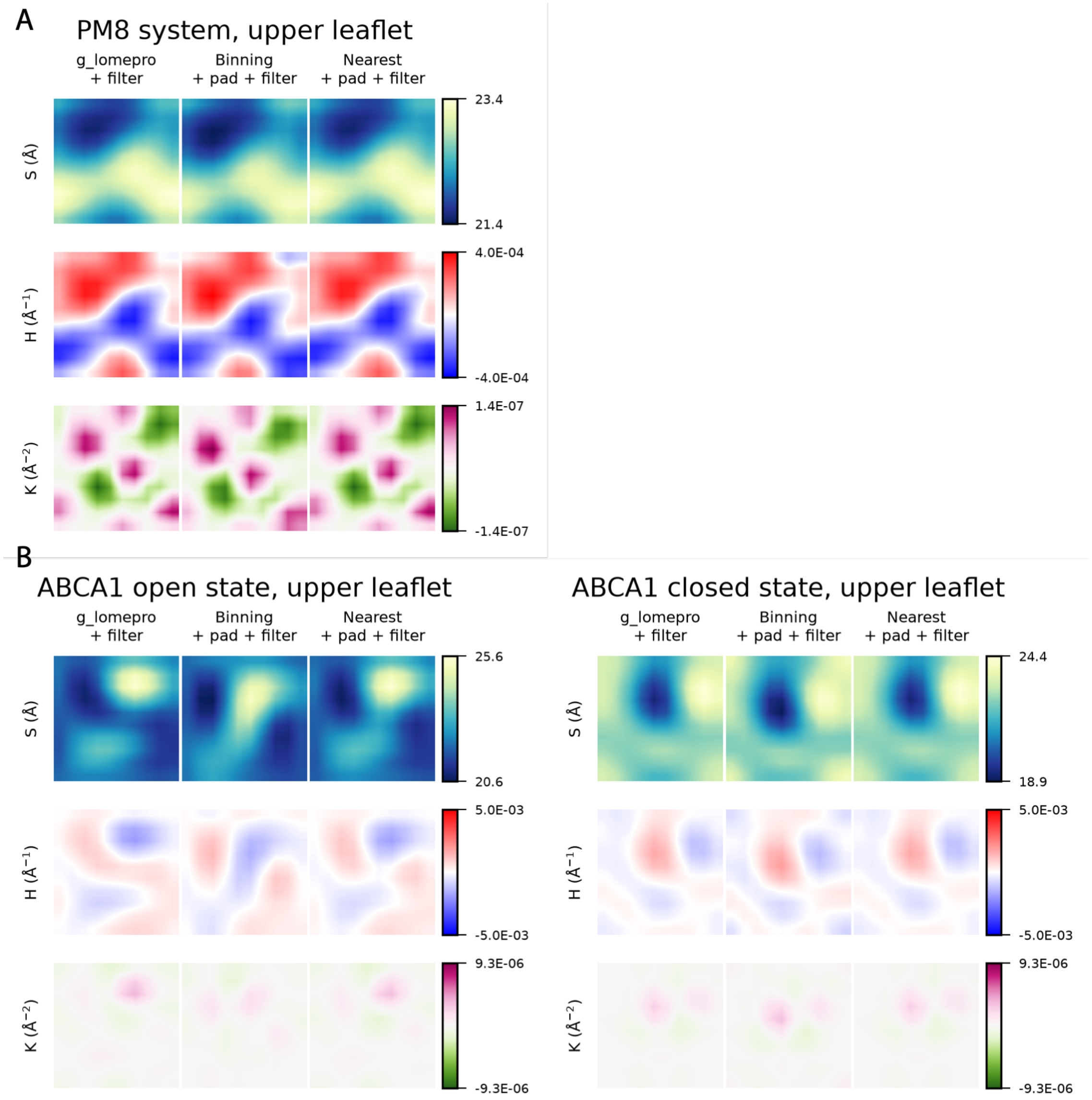


**Figure S3.** ***MembraneCurvature*** **binning methods and the FFT filter.**

The surface, Mean and Gaussian curvature of the upper leaflet of the PM8 (A) and ABCA1 systems (B) were derived for the last 5 µs of the simulations using `g_lomepro` and the binning methods in *MembraneCurvature* while applying a FFT filter, periodic padding and calculating curvature on the average surface.

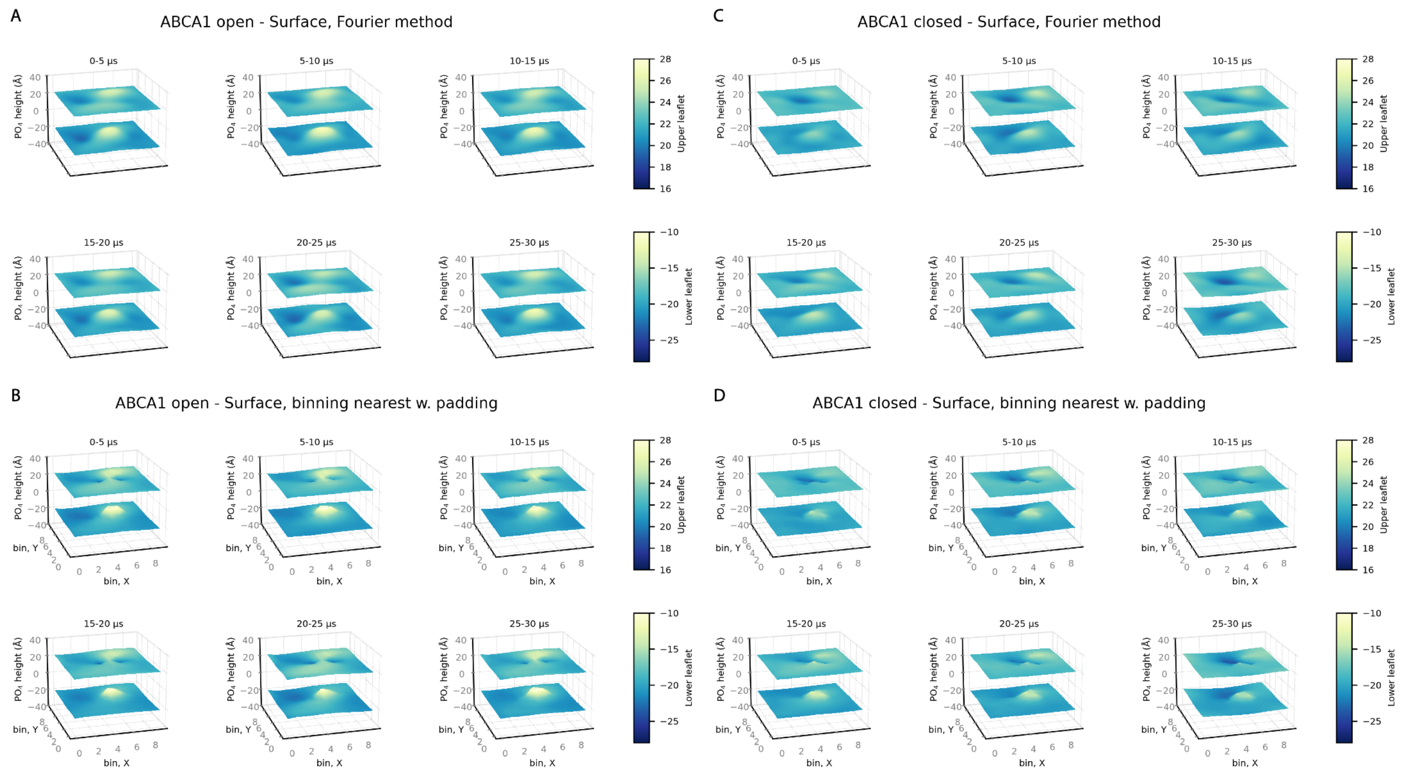


**Figure S4. Surface derivation for the ABCA1 systems using *MembraneCurvature*.**

The surface of upper and lower leaflet of the ABCA1 open (A, B) and closed (C, D) systems was derived over 5 µs-long time windows using the Fourier method (A, C), and the binning nearest method, with periodic padding applied and setting the number of bins in x and y to 10 (B, D).

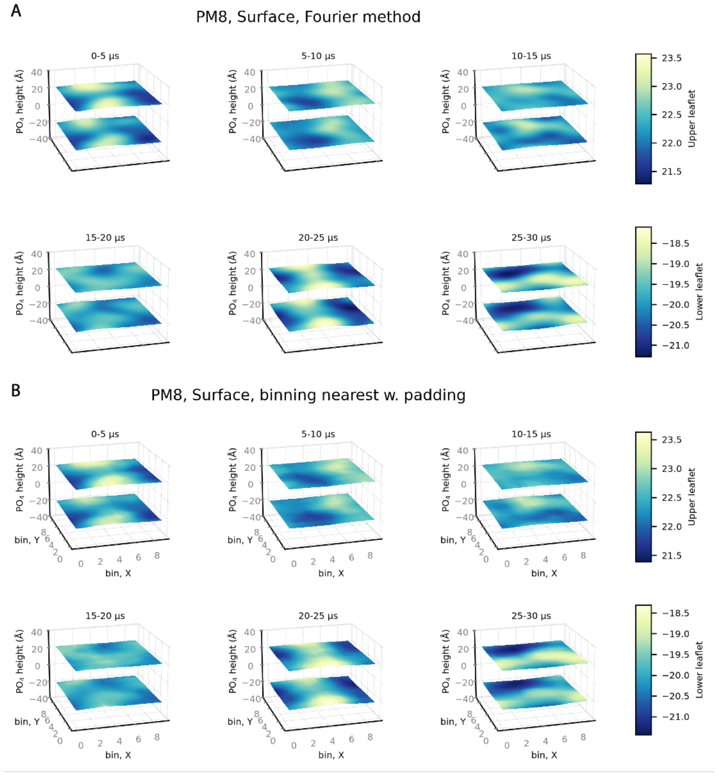


**Figure S5. Surface derivation for the PM8 system using *MembraneCurvature*.**

Surface maps derived over 5 µs time windows with *MembraneCurvature*, using the Fourier method (A) and the binning nearest method, with periodic padding applied, setting the number of bins in x and y to 10, and by using the `curvature_on='average_surface'` parameter (B).

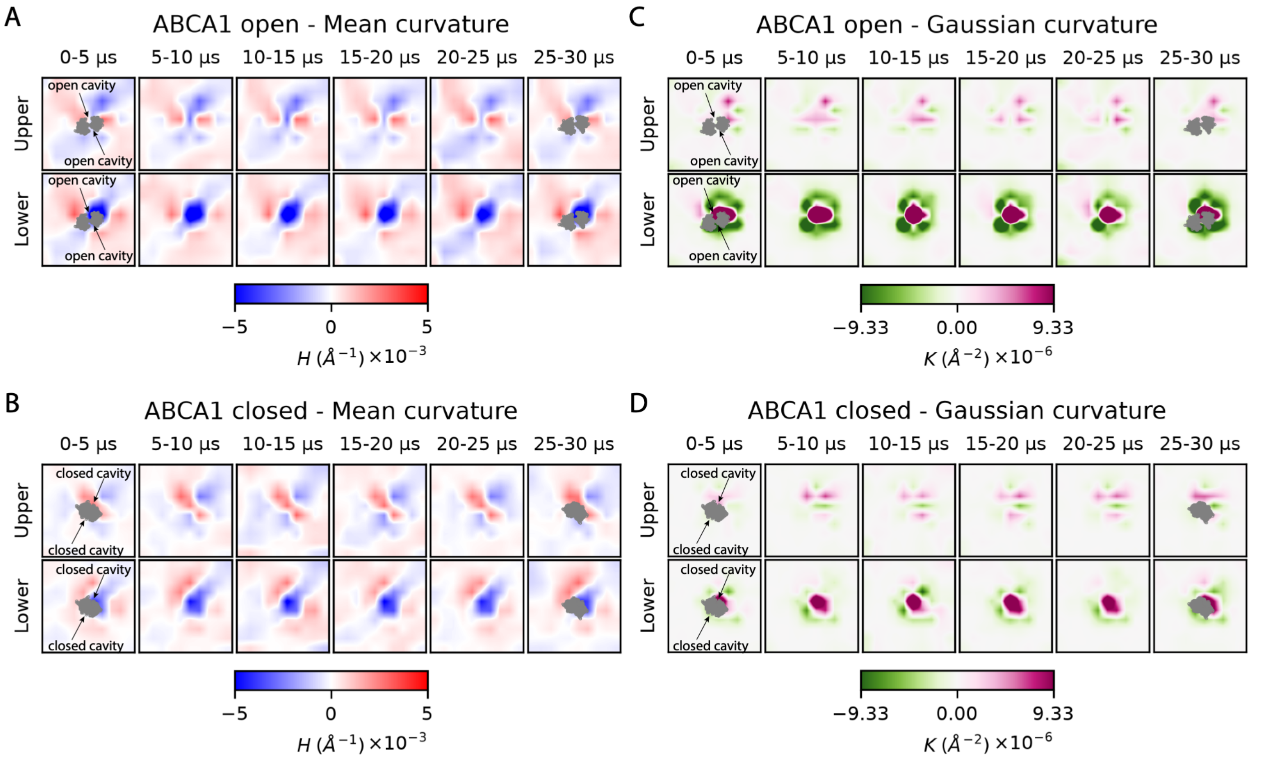


**Figure S6. Mean and Gaussian curvature maps of the ABCA1 systems calculated with the binning nearest method.**

The mean (A, B) and Gaussian (C, D) curvature maps were obtained from *MembraneCurvature* using the binning nearest method with periodic padding, setting the number of bins in x and y to 10, and by using the `curvature_on='average_surface'` parameter.

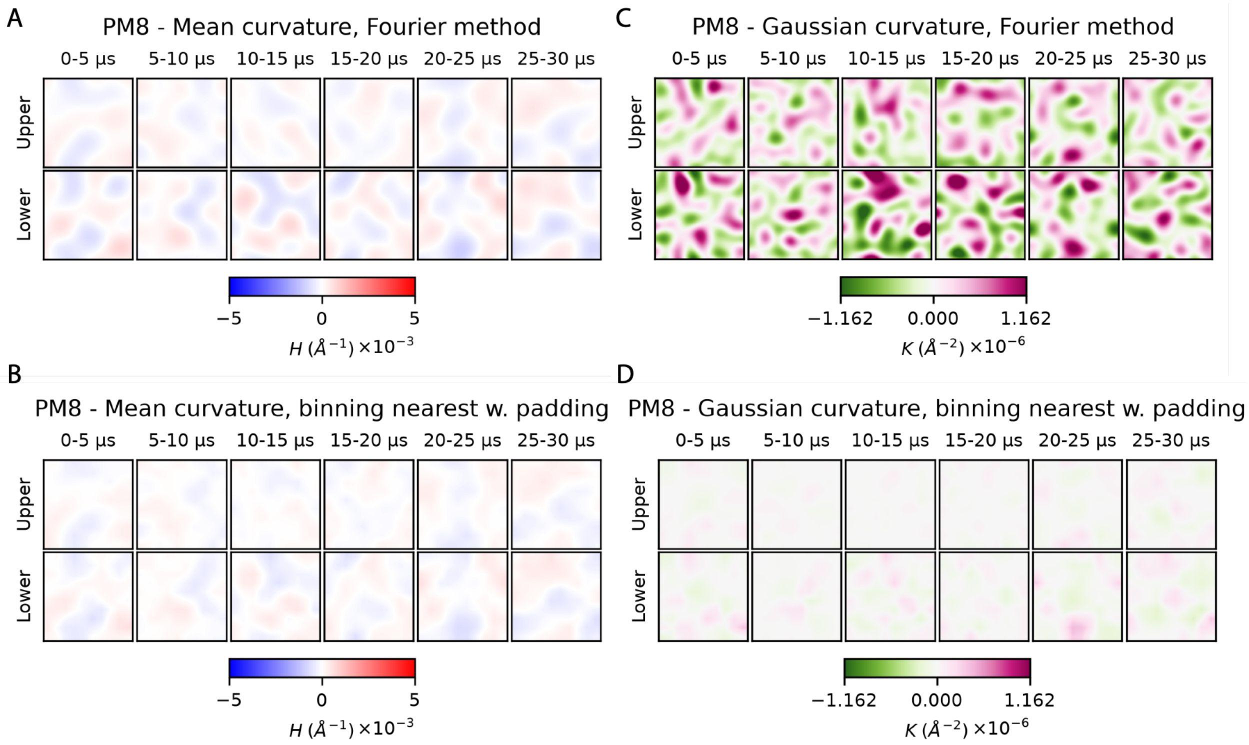


**Figure S7. Mean and Gaussian curvature maps of the PM8 system.**

The mean (A, B) and Gaussian (C, D) curvature maps were obtained from *MembraneCurvature* using the Fourier method (A, C), and the binning nearest method, with periodic padding, and setting the number of bins in x and y to 10 (B, D).

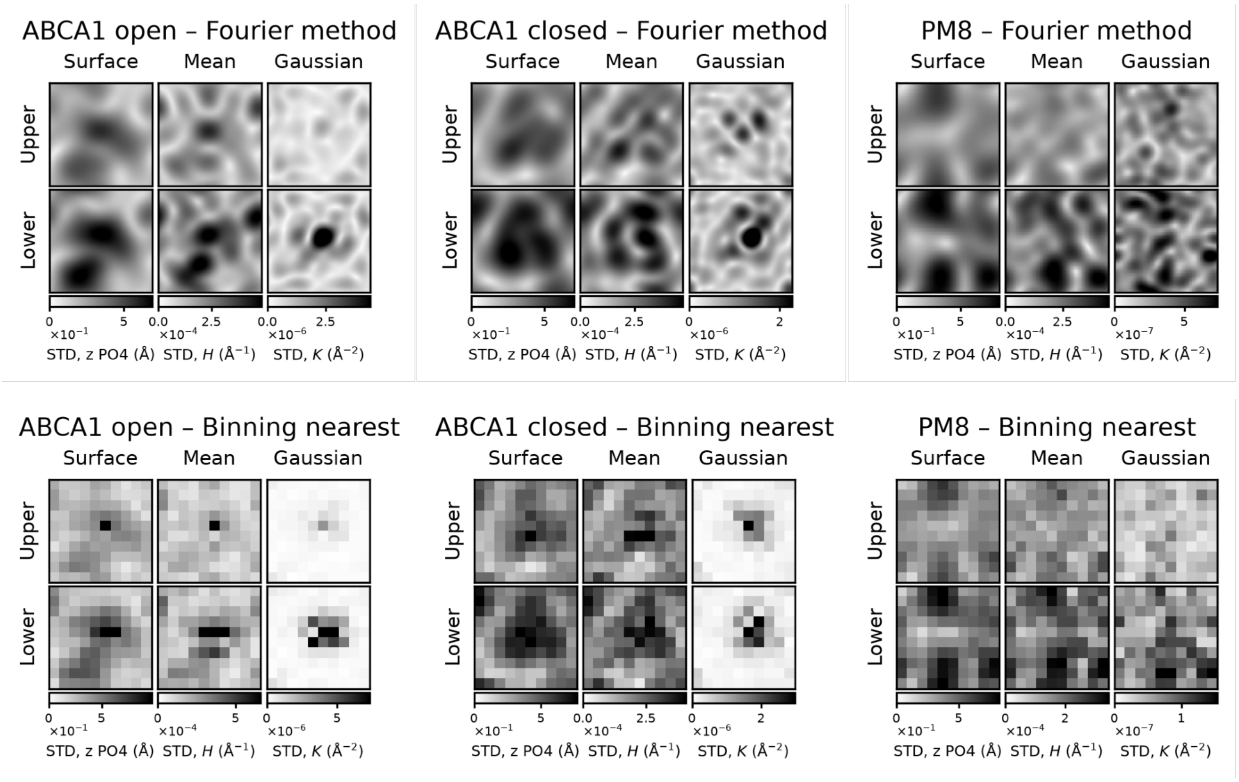


**Figure S8. Standard deviation maps for surface, mean and Gaussian curvature.**

The standard deviation maps were calculated for the surface, mean and Gaussian curvature maps obtained from the 5 μs intervals of each simulation system, using the Fourier method and the binning nearest method, with periodic padding, a 10 x 10 bin grid and with curvature calculated on the average surface (curvature_on='average_surface' parameter).

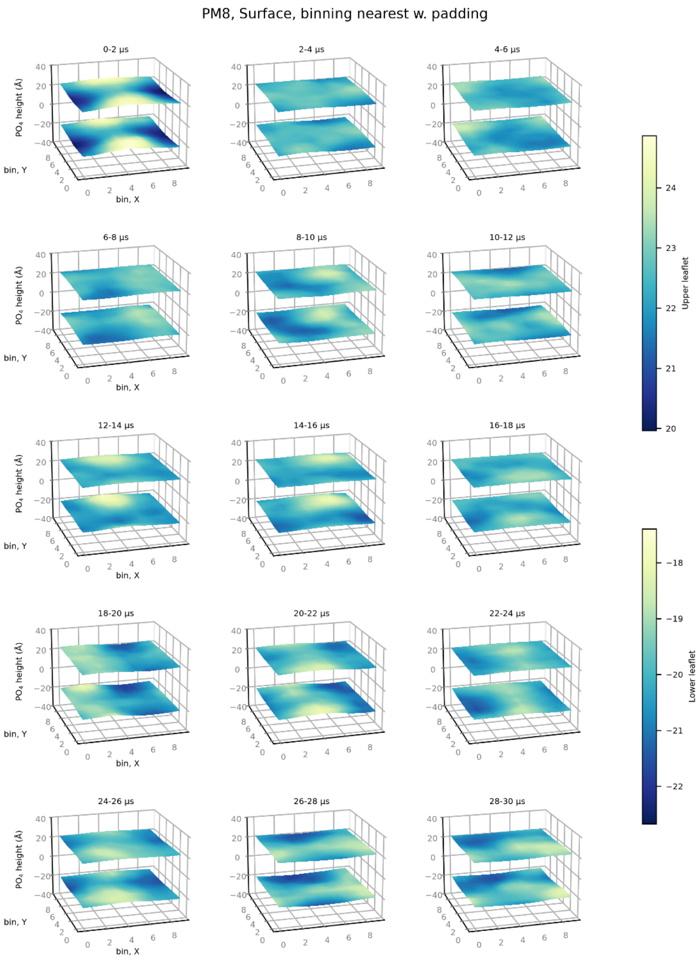


**Figure S9. Surfaces obtained from 2 μs time intervals for the PM8 system.**

The Fourier method with the default number of modes was used to derive the upper and lower leaftet surface of the PM8 system, using 2 μs time intervals spanning the 30 μs long simulation.

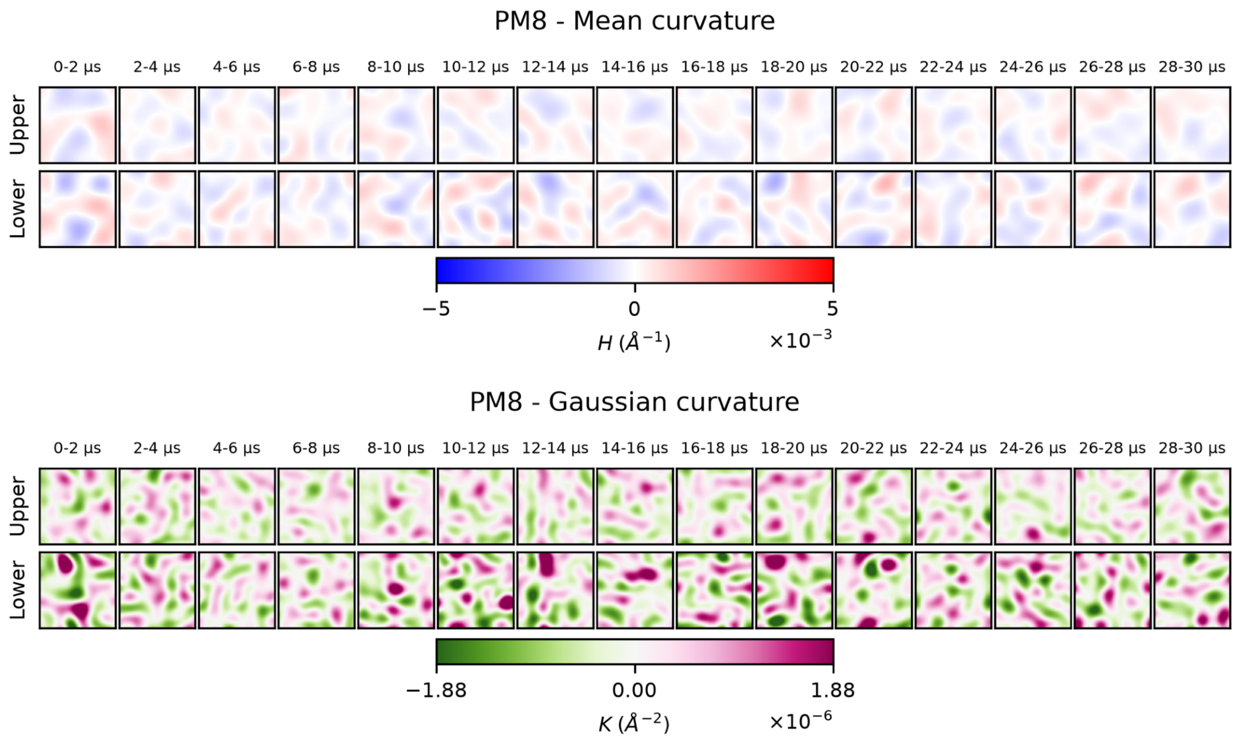


**Figure S10. Mean and Gaussian curvature from 2 µs time intervals for the PM8 system.**

The mean and Gaussian curvature maps were obtained from the corresponding surfaces of Figure S9 using the Fourier method.

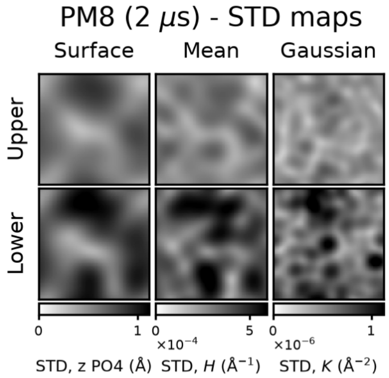


**Figure S11. Standard deviation maps for surface, mean and Gaussian curvature from 2 μs time intervals for the PM8 system.**

The mean and Gaussian curvature maps were obtained from the corresponding surfaces of Figure S10 using the Fourier method.

**Table S1. Simulation system's composition and size.**

| Molecule type | | Leaflet | # Molecules | | |
|---|---|---|---|---|---|
| | | | PM8 | ABCA1 closed | ABCA1 open |
| Lipid | POPC | Upper | 286 | 277 | 277 |
| | PAPC | | 142 | 137 | 137 |
| | POPE | | 23 | 22 | 22 |
| | DLPE | | 72 | 69 | 69 |
| | SSM | | 286 | 277 | 277 |
| | CHOL | | 414 | 400 | 400 |
| | PAPS | | 0 | 0 | 0 |
| | SAP6 | | 0 | 0 | 0 |
| | POPC | Lower | 176 | 170 | 171 |
| | PAPC | | 94 | 91 | 91 |
| | POPE | | 67 | 65 | 65 |
| | DLPE | | 205 | 198 | 199 |
| | SSM | | 137 | 132 | 133 |
| | CHOL | | 309 | 299 | 300 |
| | PAPS | | 205 | 198 | 199 |
| | SAP6 | | 28 | 27 | 27 |
| W | | | 61469 | 147000 | 147862 |
| $Na^+$ | | | 997 | 1911 | 1926 |
| $Cl^-$ | | | 680 | 1622 | 1631 |
| System size (nm)* | | | 24.57 x 24.57 x 16.49 | 24.55 x 24.55 x 34.07 | 24.63 x 24.63 x 33.95 |

*System size is reported for the final step of the 10 ns equilibration.

**Table S2. Parameters used for each surface method and curvature calculation shown in Figure 3 panels.**

| | *MembraneCurvature* parameters | | | |
|---|---|---|---|---|
| Figure 3 panel | surface_method | padding | curvature_on | fourier_m, fourier_n |
| Binning per-frame | binning | False | per_frame | – |
| Nearest per-frame | binning_nearest | False | per_frame | – |
| Binning + pad | binning | True | average_surface | – |
| Nearest + pad | binning_nearest | True | average_surface | – |
| Fourier | fourier | – | – | 2 |
